# Resolution-standardized evaluation of ligand atomic coordinates in crystallographic structures using machine learning

**DOI:** 10.64898/2026.08.17.745351

**Authors:** Ikuko Miyaguchi, Hiroaki Hata, Takaaki Kuribayashi, Shouta Takahashi, Akiko Kashima, Kouta Murasaki, Shigeyuki Matsumoto, Kei Terayama, Masateru Ohta, Mitsunori Ikeguchi

## Abstract

Accurate assessment of ligand coordinate–density consistency across different resolutions remains challenging in macromolecular crystallography. We introduce the atomic Box Correlation Coefficient (aBCC), an atom-level metric for evaluating the consistency between ligand atomic coordinates and electron density in a resolution-standardized framework. To predict aBCC values from electron-density maps, we developed QAEmap, a machine-learning model based on three-dimensional convolutional neural networks (3D-CNNs). The model was trained on ligand coordinates and Fourier-truncated electron-density maps derived from high-resolution PDB structures, using aBCC values defined against standardized 1.5 Å reference maps as training targets. It was evaluated using both Fourier-truncated electron-density maps and experimentally determined PDB structures. The prediction accuracy gradually decreased with decreasing resolution, but remained reliable up to ∼3.5 Å. These results demonstrate that aBCCpred provides a resolution-standardized atom-wise evaluation of coordinate–density consistency across different resolutions and establish a foundation for further development and refinement of machine learning-based coordinate validation.

**Synopsis:** We introduce the atomic box correlation coefficient (aBCC), an atom-level metric for the resolution-standardized atom-level evaluation of ligand coordinate–density consistency in crystallographic structures. aBCC provides a common framework for assessing and communicating the local coordinate reliability between structural biologists and researchers in structure-based drug discovery.

## 1. Introduction

The atomic coordinates of macromolecular structures deposited in the Protein Data Bank (PDB) (Berman, 2000) underpin modern structural biology, computational modeling, and structure-based drug discovery. As the number and diversity of deposited structures grow, their users increasingly include researchers who are not directly involved in crystallographic refinement (Bijak *et al.*, 2023). Consequently, intuitive and quantitative indicators of local coordinate reliability, particularly in the ligand-binding regions, are in high demand.

In current crystallographic practice, ligand placement is guided by iterative model building and refinement, and its reliability is primarily assessed through visual inspection of the fit to the electron density, complemented by validation metrics and chemical plausibility checks. However, these assessments remain partly subjective and depend on data quality (Pozharski *et al.*, 2017). Nevertheless, accurate ligand placement remains among the most challenging aspects of model building. Ligand density is frequently weakened by limited resolution, high atomic displacement, partial occupancy, and conformational heterogeneity, indicating that the local density quality is governed by multiple factors beyond the nominal resolution (Chakraborti *et al.*, 2021; Weichenberger *et al.*, 2013). Consistent with these challenges, systematic surveys have revealed that even structures nominally classified as high-resolution (≤2.5 Å) may contain ligand models that are poorly supported by electron density (Rupp *et al.*, 2016; Mir *et al.*, 2009; Gao *et al.*, 2023; Weiss *et al.*, 2022; Pozharski *et al.*, 2017).

Several validation metrics have been proposed to assess the reliability of ligand models. Among these, the real-space correlation coefficient (RSCC) is the most widely used measure of agreement between a ligand model and the observed electron density (Tickle, 2012). At the atomic level, the electron density score for individual atoms (EDIA) evaluates the local support of individual atomic positions based on electron density and has been widely adopted for assessing local model quality (Meyder *et al.*, 2017). Other metrics, including B-factors, local ligand density fit (LLDF) (Smart *et al.*, 2018), and ligand-specific scoring schemes, such as The Validation HElper for LIgands and Binding Sites (VJELIBS) (Cereto-Massagué *et al.*, 2013; Chakraborti *et al.*, 2021), provide additional perspectives on model quality.

Despite their utility, these metrics have important limitations. RSCC and EDIA are influenced by resolution, complicating direct comparisons across datasets. B-factors are affected by the refinement protocol and global model parameters, whereas LLDF is less intuitive for interpreting individual atomic positions. Ligand-specific scoring schemes, such as VJELIBS, are generally designed for the statistical analysis of large datasets rather than for the detailed evaluation of individual ligand models.

Model bias further complicates ligand evaluation (Hodel *et al.*, 1992; Terwilliger *et al.*, 2008). Because refinement procedures use an atomic model to compute phases, incorrect coordinates can reinforce erroneous density features. Although expert crystallographers address the issue by routinely inspecting Fo–Fc or polder omit maps (Liebschner *et al.*, 2017), manual assessment is not scalable and remains inaccessible to many non-specialist users. Consequently, an objective, interpretable, and resolution-standardized metric for evaluating local coordinate–density consistency is urgently needed.

Previously, we developed Quality Assessment using Electron density maps (QAEmap), a machine learning method for predicting the consistency between electron density and coordinates at the amino-acid-residue level (Miyaguchi *et al.*, 2021). QAEmap was designed to evaluate local structural quality on a standardized scale across datasets of different resolutions. However, because the original QAEmap evaluates structures at the residue level, it is unsuitable for assessing ligand molecules whose chemical identity and biological activity depend on the precise positioning of individual atoms. Small differences in orientation, torsion, or substituent placement may determine binding mode and function, and such fine-grained distinctions cannot be captured by residue-averaged metrics.

To overcome this limitation, we extended the QAEmap framework to the atomic level. We first defined the atomic Box Correlation Coefficient (aBCC), which quantifies, for each ligand atom, the local agreement between an omit electron-density map and a standardized high-resolution reference map calculated from the atomic coordinates (Fig. 1a). We then trained an atomic-level QAEmap model to predict aBCC values from electron-density maps and ligand coordinates. Fourier-truncated maps spanning a range of effective resolutions were paired with aBCC values determined from the corresponding 1.5 Å reference data. This design enables the model to learn the relationship between information loss introduced by the resolution cutoff and coordinate–density consistency evaluated against a common high-resolution reference.

**Figure 1.**
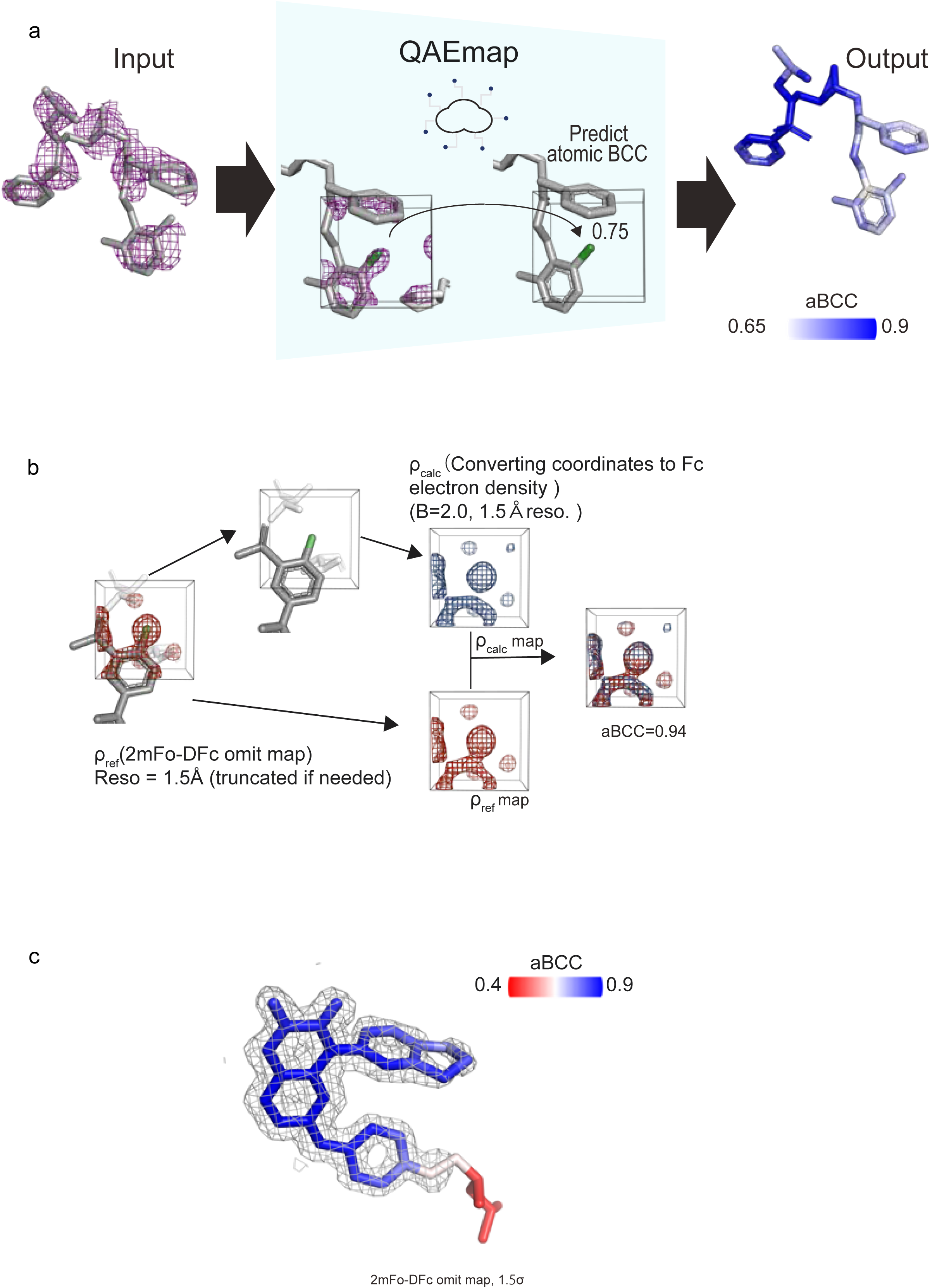
Overview of aBCC-based ligand coordinate–density consistency (a) QAEmap evaluates ligand atomic coordinates by predicting atom-wise aBCC values defined within a standardized 1.5 Å reference framework. The model was trained using data derived from high-resolution structures and predicts BCC values independently for each ligand atom. (b) aBCC is defined as the Pearson correlation coefficient computed within a cubic region (6 × 6 × 6 Å) centered on the target atom between the 2 mFo–DFc omit map and a theoretical Fc map derived from the atomic coordinates. The omit map is generated at 1.5 Å resolution (high-resolution maps are truncated as necessary), and the Fc map is calculated at 1.5 Å resolution with a fixed isotropic B-factor of 2.0 Å². In the example shown, the refined (correct) structure yielded aBCC = 0.94. (c) Interpretation of BCC values. Even after refinement, atoms with high B-factors (i.e., poorly defined positions) tended to exhibit lower aBCC values, reflecting reduced local coordinate reliability rather than refinement quality.

In this study, we establish this atomic-level QAEmap framework as a physically interpretable approach for resolution-standardized evaluation of local ligand coordinate–density consistency. The predicted aBCC values (aBCC_pred_) provide complementary information to existing validation metrics and enable individual ligand atoms to be evaluated on a common scale across a range of resolutions. The method therefore offers a practical tool for identifying locally uncertain ligand coordinates and supporting ligand-focused structural assessment in macromolecular crystallography.

## 2. Methods

### 2.1. Overview of QAEmap

QAEmap is a machine learning-based framework designed to evaluate the consistency between atomic coordinates and experimental electron density maps at the atomic level. This method defines an atom-level consistency metric, aBCC, computed between a standardized experimental reference density and a calculated density derived from atomic coordinates within a local 3D box surrounding each atom. Both the reference and calculated densities are standardized to a common resolution of 1.5 Å. A 3D convolutional neural network is trained to predict this standardized aBCC value from local density features, thereby enabling a consistent comparison of local coordinate–density consistency across structures with different resolutions (Fig. 1a).

#### 2.1.1. Input and Output

The inputs to QAEmap consist of (i) atomic coordinates derived from macromolecular crystal structures and (ii) the corresponding omit electron density maps (2 mFo–DFc) generated from the structure-factor data used for model refinement. For each atom of interest, a local cubic density box (6 × 6 × 6 Å) centered on the atomic position is extracted from the electron density map. The output of QAEmap is the predicted aBCC (aBCC_pred_), which reflects the degree of consistency between the atomic model and experimental electron density on a per-atom basis.

#### 2.1.2. Definition of aBCC

aBCC is defined as the Pearson correlation coefficient between the reference and calculated electron density values within a local cubic box surrounding an atom (Fig. 1b).

The reference density (ρ_ref_) was obtained from omit maps (2 mFo–DFc) generated by removing the target ligand prior to phase calculation, thereby minimizing model bias. To standardize the density across structures, the omit maps were Fourier-truncated to 1.5 Å resolution regardless of the experimental resolution. When higher-resolution data were available, the structure factors were truncated.

The calculated density (ρ_calc_) was obtained from theoretical Fc maps computed from the atomic coordinates with all atomic B-factors fixed to an isotropic value of 2.0 Å² and using structure factors truncated to 1.5 Å resolution, ensuring that the density reflects only coordinate-dependent features rather than variations in atomic displacement parameters. For each atom, density values were sampled on a regular grid within a cubic box (6 × 6 × 6 Å) centered at the atomic coordinates. Both reference and calculated maps were sampled at a grid spacing of 0.3 Å, which preserves fine structural features observable at 1.5 Å resolution.

The aBCC is defined as:

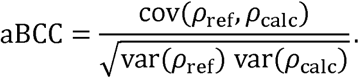

Atoms with well-defined coordinates and clear supporting density yield high aBCC values approaching 1.0, whereas atoms associated with weak or ambiguous density—arising from positional disorder, high mobility, or incorrect placement—exhibit lower aBCC values (Fig. 1c, Supplementary Fig. 1). Importantly, a low aBCC value does not necessarily indicate incorrect atomic coordinates; rather, it indicates weak local agreement between the atomic model and the experimental electron density. The use of a 6 Å box allows the metric to incorporate not only the density at the target atom but also the surrounding chemical environment.

The resulting values were used as reference values in this study and are referred to as aBCC_act_ (actual aBCC values).

### 2.2. Construction of training data for machine learning model

#### 2.2.1. Fourier-truncated maps and training data

The basic concept underlying the construction of the training data was to relate the information lost through Fourier truncation of high-resolution diffraction data to coordinate quality evaluated against a standardized 1.5 Å reference. The training data were therefore designed to allow the effect of resolution-dependent information loss to be learned separately from density variations caused by other factors, including resolution-dependent changes in overall B factors. The Fourier-truncated maps used in this study were not intended to reproduce all characteristics of experimental datasets collected at lower resolutions. Instead, they provided systematically controlled inputs in which the loss of high-resolution information was introduced while the coordinate models were kept unchanged across resolution levels.

To construct these training data, high-resolution datasets with an experimental resolution of 1.5 Å or better were selected. Atomic coordinates were obtained from deposited PDB models after removing water molecules. Only atoms with an occupancy of 1.0 and without alternative conformations were included.

The training data comprised atomic coordinates and the corresponding omit electron-density maps. To systematically vary the information available in the input maps while preserving other properties of the datasets, structure-factor amplitudes were truncated in 0.5 Å increments from 2.0 to 4.0 Å. The corresponding 2mFo–DFc omit maps were generated using the CCP4 program suite. For each structure, the same coordinate models, including their atomic B factors refined against data truncated to 1.5 Å, were retained at all resolution levels. Thus, only the resolution cutoff of the input map was varied, whereas the corresponding aBCC_act_ value remained identical across resolutions (Supplementary Fig. 3).

Although atomic B factors were not varied as a function of resolution, the training coordinate models retained substantial variation in ligand B factors, ranging from 3 to 74 Å², with mean ligand B factors ranging from 7 to 61 Å² among the models (Supplementary Table 2). For model input, local density features were extracted for each atom using cubic boxes (6 × 6 × 6 Å) centered at the atomic coordinates. Electron density values were sampled on a regular grid with a spacing of 0.3 Å, consistent with the aBCC calculation. The same spatial region was used to extract density values from omit maps at each resolution level. Atomic identity was encoded using atom type categories (CONS), with all other atoms grouped into a single category (X) (Fig. 2a and Supplementary Fig. 4). The procedures for local feature extraction and pre-processing followed those described in our previous study (Miyaguchi *et al.*, 2021).

**Figure 2.**
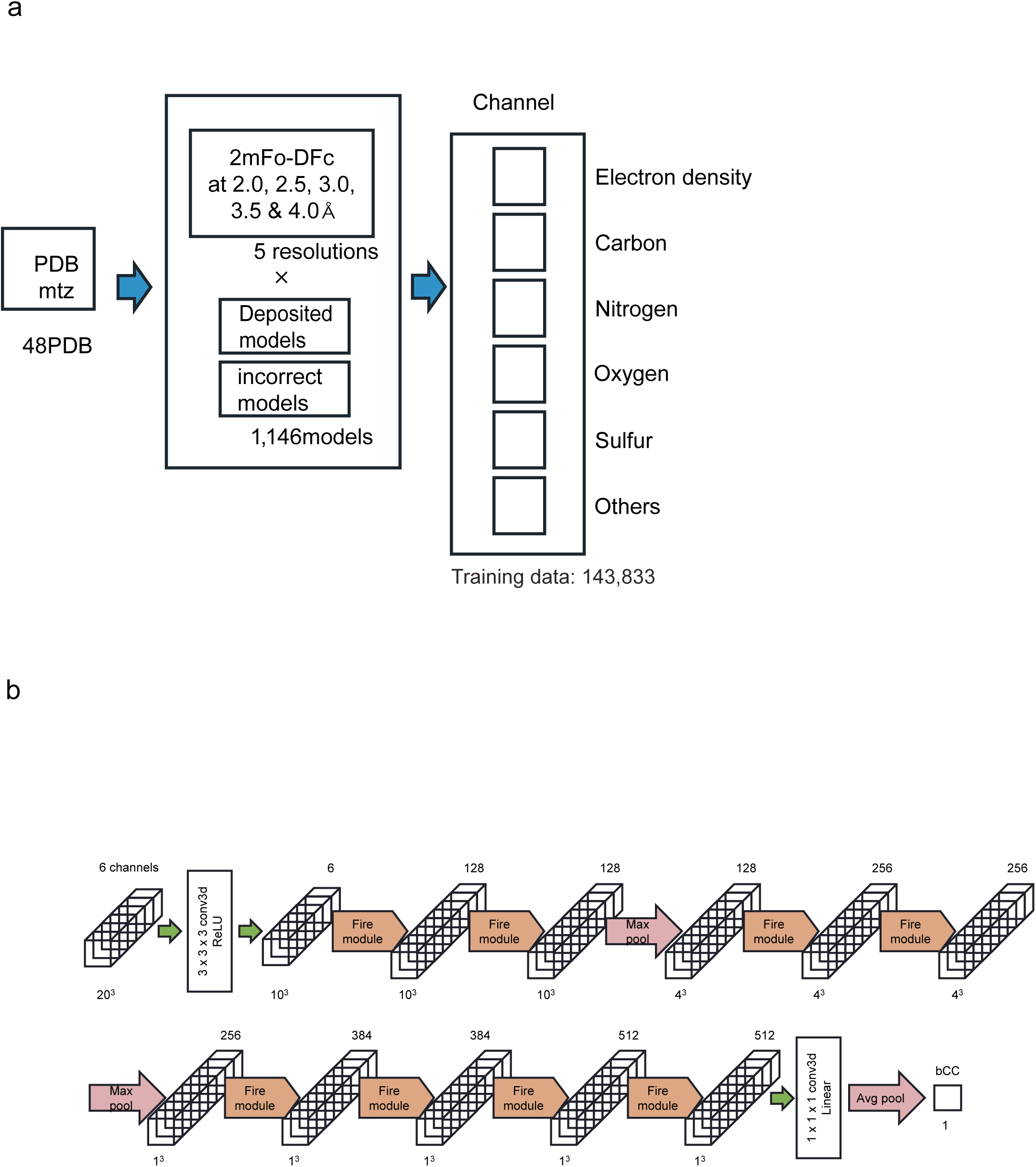
Training data preparation and model architecture of QAEmap. (a) Workflow for generating training data from the PDB structures. Starting with 48 entries, 2 mFo–DFc omit maps were generated at five resolutions (2.0, 2.5, 3.0, 3.5, and 4.0 Å). For each structure, both the deposited and incorrect models were used to capture diverse atomic environments, yielding 1,400 models. For each atom, six-channel input volumes were constructed comprising one electron-density channel and five element-type channels (C, N, O, S, and others). (b) Model architecture. The network was based on a SqueezeNet-style 3D convolutional architecture, beginning with a 3 × 3 × 3 convolutional layer applied to six-channel input volumes representing local atomic environments. This is followed by multiple Fire modules, each comprising a squeeze layer and expanded layers interleaved with max-pooling layers for hierarchical feature extraction. The network was trained as a regression model to predict the aBCC_pred_.

The test datasets used for performance evaluation were prepared using the same protocol and comprised 12 protein–ligand pairs spanning multiple resolution levels. Training data were generated using Fourier-truncated maps at 2.0–4.0 Å resolutions, corresponding to the intended operating range of the model. Consequently, the 1.5 Å maps were excluded from the training dataset and reserved for independent evaluation.

The structures were obtained from the PDB. Sixty entries were used to construct the dataset, of which 12 were reserved as independent test sets. The number of models and data points used for training are summarized in Table 1, and the details of the compounds included in the training dataset are provided in Supplementary Tables 1 and 2. The structural diversity of the compounds used for training was assessed using principal component analysis (PCA) and uniform manifold approximation and projection (UMAP) based on molecular descriptors (Supplementary Fig. 5). The results suggested that the selected compounds were distributed across a relatively broad chemical space.

**Table 1.** Summary of training data.

|  |  | Training data | Test data |
|---|---|---|---|
| PDB |  | 48 | 12 |
| Models |  | 5730 | 1267 |
| Data |  | 143833 | 30372 |
| Atom | C | 95556 | 20182 |
|  | N | 16062 | 4896 |
|  | O | 28621 | 4668 |
|  | S | 1161 | 313 |
|  | P | 1590 | 106 |
|  | F | 424 | 0 |
|  | Cl | 313 | 207 |
|  | Br | 106 | 0 |

#### 2.2.2. 3D convolutional neural network architecture

The model used to predict aBCC followed a 3D SqueezeNet-style architecture and was implemented according to our previous protocol (Miyaguchi *et al.*, 2021). The network begins with an initial 3D convolutional layer, followed by a stack of Fire modules (each comprising squeeze and expand operations) interleaved with max-pooling layers (Fig. 2b). After the final Fire module, a dropout layer (rate = 0.5) was applied, and a (1 × 1 × 1) Conv3D layer was used to reduce the channel dimensionality, followed by global average pooling; the pooled scalar is returned as the single regression output, aBCC_pred_. The loss function was the mean squared error (MSE).

The input representation and preprocessing steps are as follows: Local 3D regions are encoded as cubic boxes of physical size (6 × 6 × 6 Å) sampled on a 0.3 Å grid, producing voxel volumes of dimension 20^3^. Each input voxel has six channels corresponding to atom types C, N, O, S, and X (other) and the electron density map. During pre-processing, each box is augmented by rotations in 90° increments, yielding 24 orientations per sample.

Training details: The models were trained using a hold-out strategy in which the full dataset was first split into training and test sets at an 80/20 ratio. The training portion was further divided into training and validation subsets using an additional 80/20 split. Training was performed with a batch size of 64 and a maximum of 100 epochs using the Adam optimizer with a learning rate of 1 × 10^-3^ and MSE as the objective function. Early stopping with a patience of 10 epochs was performed to prevent overfitting. These implementation and training choices followed the previously established workflow (Miyaguchi *et al.*, 2021) to maximize reproducibility and facilitate a direct comparison with earlier results.

### 2.3. Rationale for the standardized reference density used in aBCC (1.5 Å, B = 2.0 Å²)

We next describe the rationale for adopting standardized conditions for calculating reference electron density, namely a resolution cutoff of 1.5 Å with a fixed B-factor of 2.0 Å².

The aBCC evaluates the consistency between atomic coordinates and high-resolution reference electron density based on the premise that coordinate reliability is best assessed when individual atomic features are explicitly resolved. In practice, resolutions around 1.5 Å represent a widely accepted threshold at which atomic density peaks are still clearly separated and interpretable.

Although even higher resolution cutoffs can, in principle, provide more detailed information, their practical utility is limited by the strong attenuation of high-frequency Fourier components caused by B-factors (Fig. 3a). This attenuation substantially reduces the contribution of the high-resolution terms, diminishing the effective gain from extending the nominal resolution beyond ∼1.5 Å.

**Figure 3.**
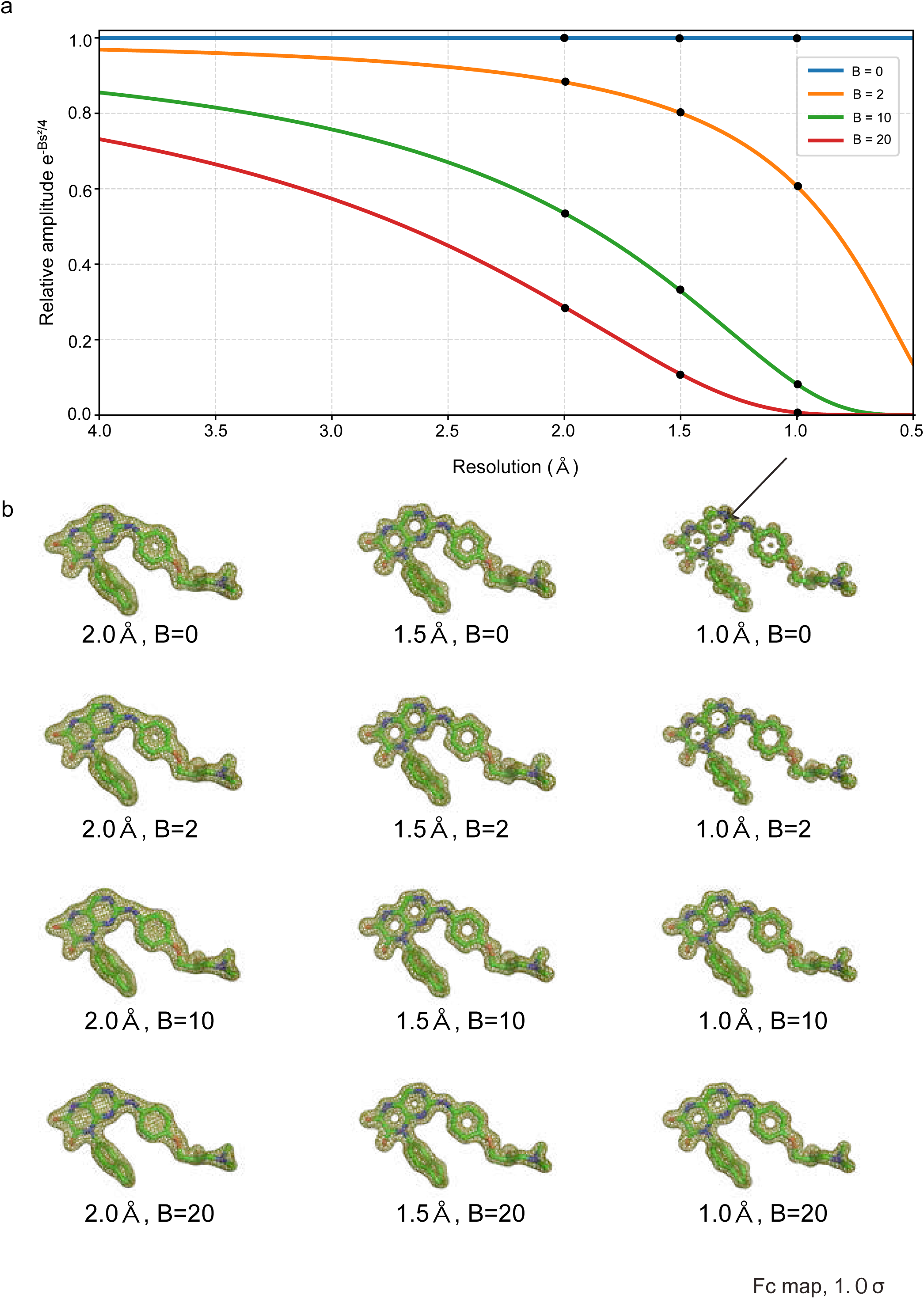
Rationale for defining the standardized reference map (1.5 Å, B = 2.0 Å²). (a) Attenuation of the structure-factor amplitudes as a function of resolution for different B-factors. Increasing the B-factors leads to stronger damping of the high-resolution Fourier components, limiting the effective contribution of the high-resolution terms even when the nominal resolution is extended. (b) Examples of Fc maps calculated at selected points indicated in panel (a) using the ligand KLM from PDB entry 6NFH. Maps calculated with low B-factors (e.g., B = 0 Å²) at high nominal resolution exhibit pronounced truncation artifacts, including Gibbs oscillations (arrow). In contrast, a small and finite B-factor (e.g., B = 2.0 Å²) suppresses these artifacts while preserving interpretable atomic features.

At lower resolutions (e.g., ∼2.0 Å), electron density increasingly merges into continuous features and loses atom-wise discriminability, rendering such maps suboptimal for evaluating coordinate accuracy at the atomic level (Fig. 3b). Conversely, setting the B-factors to zero would fully preserve the high-resolution amplitudes; however, this produces electron density distributions that deviate markedly from the experimental reality and introduce pronounced Fourier truncation artifacts, including Gibbs oscillations. These non-physical features generate spurious density peaks and inflate apparent model–map discrepancies (Fig. 3b, B = 0 at 1.0 Å; Supplementary Fig. 6, B = 0 at 1.5 Å), rendering B = 0 inappropriate for defining a reference density.

We therefore adopted a small fixed B-factor (B = 2.0 Å²) in combination with a 1.5 Å resolution cutoff when calculating reference Fc maps. This choice represents a practical compromise: it preserves sufficient high-resolution information for atom-wise evaluation while suppressing unphysical oscillations and excessive deviation from realistic electron density distributions. Importantly, this condition is not intended to reproduce experimental B-values but to provide a standardized, sharply localized high-resolution reference. Using experimental B-factors in the reference map could obscure insufficient experimental support for poorly ordered atoms, because their broadened density would contribute less to the correlation, while better-ordered neighboring atoms within the 6 Å box could dominate it.

The uniformly low B-factor ensures that poor local experimental support results in a reduced aBCC, rather than being masked by a corresponding broadening of the calculated Fc density.

In addition to defining the reference density using standardized Fc maps (1.5 Å, B = 2.0 Å²), omit maps (2mFo–DFc with the target atoms removed) calculated at the same resolution cutoff were used for aBCC evaluation to minimize model bias. This approach reduces artificial model–map consistency arising from refinement procedures.

Together with element-specific input channels, this framework is designed to assess local, atom-wise coordinate–density consistency against a common reference. Consequently, aBCC is expected to reduce systematic resolution-dependent variation while retaining sensitivity to genuine local disorder reflected in the experimental density.

### 2.4. Dataset preparation and evaluation procedures

#### 2.4.1. Performance evaluation datasets

To evaluate the performance of the aBCC model, two types of datasets were analyzed.

First, the held-out test dataset defined during model development was used to assess the predictive accuracy under controlled conditions. For this dataset, the aBCC_act_ values were calculated using the standardized reference map conditions, enabling a direct comparison between the predicted and reference values.

Secondly, an independent real-structure dataset (PDB Ligand ID: 23E) comprising experimentally determined ligand-bound protein structures that were not included in the training or test sets was analyzed to examine the behavior of the model in practical applications. These structures were selected from the PDB group depositions and filtered to include only ligand-bound entries while retaining only those with the same crystal form within the group depositions. The aBCC predictions (aBCC_pred_) were computed directly from the deposited models without calculating the reference aBCC values (aBCC_act_).

For each structure, water molecules were removed, and the atomic coordinates were used without modification. Omit maps were used for the aBCC prediction. A complete list of the datasets analyzed in this study is provided in Supplementary Table 3.

#### 2.4.2. Experimental omit-map aBCC_act_ and EDIA calculations

Experimental omit-map aBCC_act_ values were calculated using the same procedure as that used for the aBCC_act_ calculation during training data preparation, except that experimentally derived omit maps were used instead of artificially resolution-truncated omit maps. Atomic EDIA values were obtained using the EDIA server (Meyder *et al.*, 2017). For each structure included in the multiresolution 23E dataset, the EDIA scores were retrieved by specifying the corresponding PDB entries. The experimental omit-map aBCC_act_ and atomic EDIA values were compared across structures of different resolutions and evaluated alongside the atomic aBCC_pred_ values.

#### 2.4.3. SAM, FMN, and ATP datasets and RSCC calculation

To assess the resolution dependence of aBCC_pred_ compared with conventional real-space correlation metrics, structures containing the ubiquitous cofactors S-adenosylmethionine (SAM), flavin mononucleotide (FMN), and adenosine triphosphate (ATP) were systematically collected from the PDB. These cofactors are frequently observed in protein structures and are represented by numerous entries across a broad resolution range.

Only ligands with full occupancies (occupancy = 1.0) and complete atomic models were included in the analysis. Entries containing partially modeled ligands or missing atoms were excluded. In addition, structures for which the aBCC could not be calculated (e.g., owing to map availability or formatting inconsistencies) were removed from the dataset.

The curated structures were grouped into resolution bins, and the aBCC_pred_ values were calculated for each ligand directly from the deposited models and corresponding omit maps. In parallel, ligand-level RSCCs were calculated independently using Phenix based on conventional 2 mFo–DFc maps. RSCC calculations were performed only for structures for which the aBCC values were successfully obtained.

In some cases, the RSCC calculation failed for technical reasons related to map generation or formatting. Such entries were excluded from analyses involving RSCC, whereas their aBCC_pred_ values were retained in the aBCC dataset. For direct comparison between the two metrics, only the subset of structures for which both aBCC and RSCC values were available was used. To ensure fair comparison with aBCC_pred_, the map grid spacing was set to 0.3 Å for RSCC calculations. Both metrics were subsequently analyzed as functions of resolution.

The complete list of SAM, FMN, and ATP datasets analyzed in this study, together with the relevant structural information, is provided in Supplementary Tables 4–6.

#### 2.4.4. Obsolete and replaced structure datasets

To examine whether aBCC_pred_ could detect changes in ligand modeling associated with structure replacement, pairs of obsolete and replaced PDB entries were analyzed. Obsolete entries and their corresponding replacement structures were obtained from the PDB archive (accessed September 2024). Only the entries containing bound ligands were considered.

Several filtering criteria were applied to ensure a meaningful comparison between the structures before and after replacement. Pairs were retained only when the ligand identity was identical between the obsolete and replacement entries and the ligand was bound to the same binding site. Structures containing buffer components or small nonspecific molecules were excluded from the analysis. Ligands with occupancies different from 1.0 were also removed to avoid ambiguity arising from partial occupancy models. In addition, pairs showing minimal structural change [ligand root-mean-square deviation (RMSD) ≤ 1.4 Å between the obsolete and replacement models] were excluded, as such cases do not represent substantial model revision. Structures for which aBCC_pred_ calculations could not be performed were also discarded.

After applying these criteria, 15 obsolete–replacement pairs of protein–ligand complexes were retained for analysis.

The aBCC_pred_ values were calculated for both obsolete and replacement structures under identical reference map conditions. The resulting scores were compared to assess whether aBCC_pred_ reflects changes in ligand–electron density consistency following structure replacement.

The complete list of analyzed obsolete–replacement pairs, together with ligand identities, resolutions, and structural details, is provided in Supplementary Table 7.

## 3. Results and Discussion

### 3.1. Performance of machine learning model

#### 3.1.1. Correlation between predicted and reference aBCC (aBCC_pred_ vs aBCC_act_)

To evaluate the performance of the trained model, we first assessed the accuracy of the aBCC prediction using independent test datasets generated in the same manner as the training data. Because the target variable was defined such that identical atomic coordinates would ideally yield the same aBCC_pred_ value across different map resolutions, accurate prediction across different resolutions would indicate a resolution-standardized coordinate evaluation.

Figure 4a shows the correlation between aBCC_pred_ and aBCC_act_ values for the test datasets at different resolutions. At 2.0 Å, the model achieved a high coefficient of determination (*R*² = 0.93), indicating that the trained model accurately reproduces the target aBCC values. Although the correlation gradually decreased as resolution decreased, a substantial correlation was maintained even at 4.0 Å (*R*² = 0.70), demonstrating the robustness of the model over a range of resolutions.

**Figure 4.**
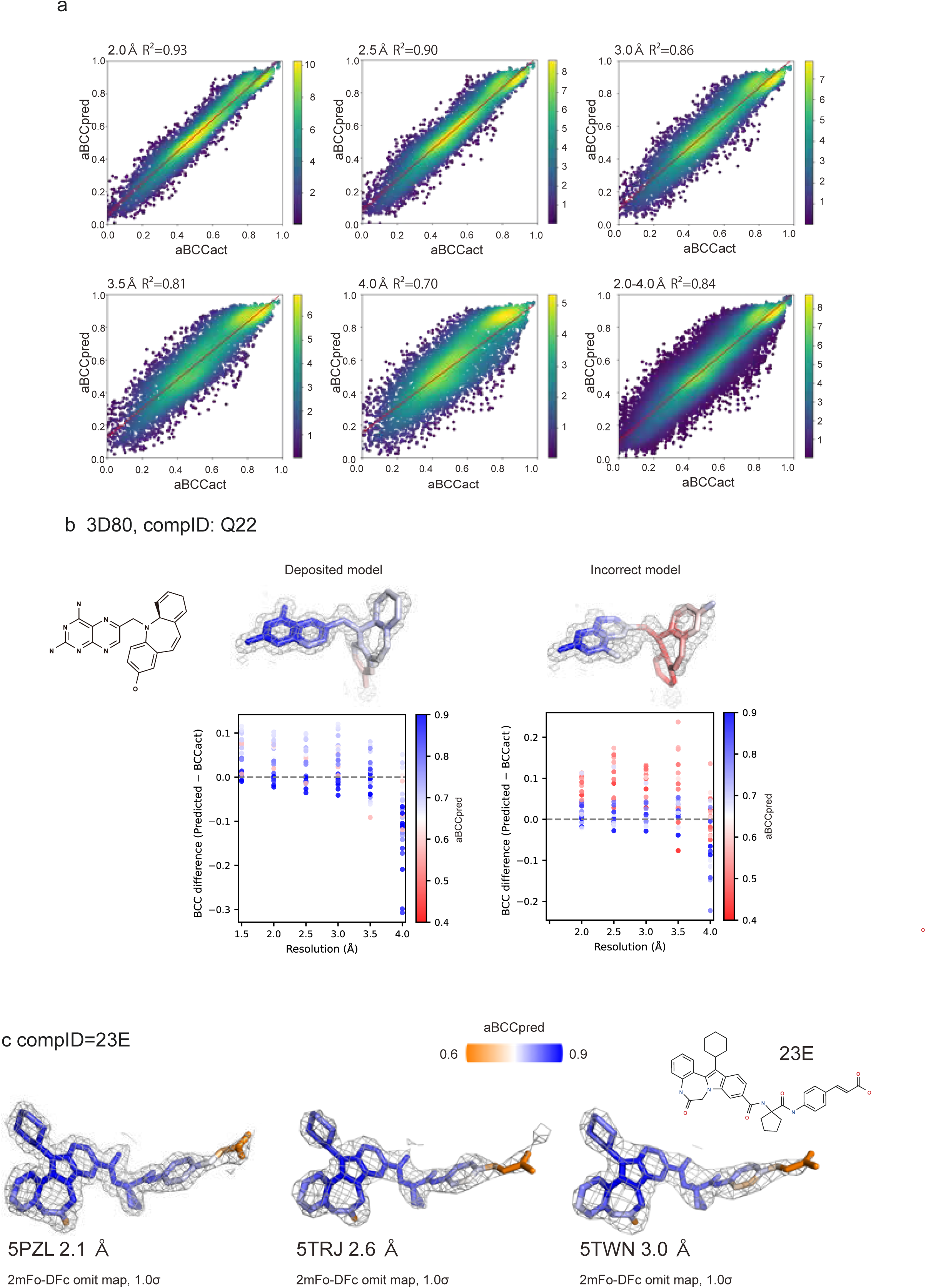
Performance of QAEmap across resolutions. (a) Correlation between reference aBCC (aBCC_act_) and predicted aBCC (aBCC_pred_) for test datasets at different resolutions (2.0, 2.5, 3.0, 3.5, and 4.0 Å), together with the pooled dataset (2.0–4.0 Å). Each point represents an individual atom. The color scale reflects the local point density estimated using kernel density estimation, with warmer colors indicating higher densities. Color scales were normalized independently for each panel. (b) Resolution-dependent behavior of aBCC prediction for representative correct and incorrect ligand models (PDB ID: 3D80). For each atom, the difference between aBCC_pred_ and the reference aBCC_act_ was plotted as a function of map resolution. The ligand models shown above the plots and the individual data points are colored according to their reference aBCC_act_ values using the red–white–blue color scale shown. The model was trained using maps at resolutions of 2.0–4.0 Å; therefore, the predictions at 1.5 Å represent extrapolation beyond the training range. (c) Representative atom-wise aBCC_pred_ distributions for ligand 23E in independently determined structures at 2.1, 2.6, and 3.0 Å resolution. Ligand atoms are colored according to their aBCC_pred_ values using the orange–white–blue color scale shown. A similar spatial pattern was observed across the three structures: the well-defined part of the ligand consistently showed relatively high aBCC_pred_ values, whereas the disordered region showed lower values regardless of resolution. This comparison illustrates that the atom-wise aBCC_pred_ pattern was largely conserved across experimental datasets of different resolutions.

#### 3.1.2. Resolution-dependent behavior of aBCC prediction on test datasets

To further examine the resolution dependence at the atomic level, we analyzed how the aBCC_pred_ values changed with the resolution for the individual ligand atoms (Fig. 4b). Using the reference aBCC_act_ values as a baseline, we evaluated the deviation of the predictions at a lower resolution. Individual atoms were color-coded according to their reference aBCC_act_ values, allowing the comparison of prediction errors across atoms with different confidence levels.

Atoms with relatively high reference values (aBCC_act_ > 0.7) exhibited small prediction errors throughout the tested resolution ranges. Up to 3.5 Å, deviations for these atoms were generally within ∼5%, indicating that well-supported atomic coordinates can be evaluated consistently across resolutions. In contrast, atoms with lower reference values exhibited larger prediction errors. In the representative correct and incorrect models shown in Fig. 4b, aBCC_pred_ tended to be higher than aBCC_act_ for these atoms, with overestimation reaching approximately 0.1 in some cases. For a given ligand model, the errors tended to exhibit consistent patterns across resolutions, although their patterns varied among the different ligand models (Supplementary Fig. 7). These observations indicate reduced quantitative accuracy in the lower range of aBCC values.

Representative examples are shown in Supplementary Fig. 7. Although the prediction errors gradually increased with decreasing resolution, the overall spatial distribution of aBCC_pred_ values remained largely unchanged. Regions predicted to have high or low coordinate reliability were consistently identified over the tested resolution range, suggesting that the model preserved local quality patterns even when the prediction accuracy decreased at lower resolutions.

Taken together with the correlation analysis in Fig. 4a, these results indicated that the trained model provides an approximately resolution-standardized assessment of the ligand atomic coordinate quality while maintaining consistent spatial quality patterns, at least up to ∼3.5 Å resolution.

#### 3.1.3. Evaluation of resolution-standardized aBCC prediction using experimental datasets

We next examined whether aBCC_pred_ provides a resolution-standardized assessment when applied to experimental crystallographic datasets. Unlike the resolution-controlled test data, which were generated by Fourier truncation of high-resolution datasets, independently collected experimental datasets contain additional variation arising from differences in overall signal attenuation, disorder, noise, and other crystal- and dataset-specific properties. We therefore performed two complementary analyses: a Fourier-truncation experiment using a single experimental dataset to isolate the effect of resolution cutoff, and a comparison of independently determined structures to evaluate the robustness of the predictions under variation present in real crystallographic data.

First, the diffraction data for 5PZL, originally determined at 2.1 Å resolution, were truncated to resolutions ranging from 2.3 to 3.0 Å, and aBCC_pred_ was recalculated using the resulting maps. Because the same diffraction dataset and coordinate model were used throughout this analysis, differences among these predictions primarily reflect the loss of high-resolution reflections. The atom-wise aBCC_pred_ profiles remained nearly unchanged after truncation, with differences of approximately 0.01 in the mean aBCC_pred_ value (Supplementary Fig. 8a, Supplementary Table 3). Based on our training-data design, we had expected aBCC_pred_ to remain relatively stable despite the loss of high-resolution Fourier information, and the results obtained with the Fourier-truncated 5PZL series were consistent with this expectation.

Having confirmed this behavior in the controlled Fourier-truncation analysis, we next examined whether aBCC_pred_ remained stable across independently collected experimental datasets differing in resolution and other dataset-specific properties. Figure 4c shows ligand 23E bound to the same protein in the same crystal form and determined at resolutions of 2.1, 2.6, and 3.0 Å. Despite visible differences in map quality, the overall aBCC_pred_ profiles were similar across these structures at both the molecular and atomic levels, although small differences remained among the datasets.

To examine this behavior more broadly, we analyzed all 13 available structures containing ligand 23E (Supplementary Fig. 8b and Supplementary Table 3). The atom-wise aBCC_pred_ profiles were generally conserved across the datasets. A modest decrease in aBCC_pred_ was observed toward lower resolution, but this dependence was limited compared with that of metrics calculated directly from the experimental maps, as described below. The variation among the independently determined structures was somewhat greater than that observed in the Fourier-truncated 5PZL series, consistent with the presence of crystal- and dataset-specific differences in addition to the effect of resolution cutoff.

To assess the reproducibility of the predictions at the atomic level, we calculated the standard deviation (SD) of aBCC_pred_ for each atom across the 13 datasets and compared it with the positional SD derived from the corresponding atomic coordinates (Supplementary Fig. 8c). Several atoms exhibited relatively large positional SDs, reflecting differences in ligand conformation among the independently refined structures. Variability in aBCC_pred_ for these atoms may therefore reflect genuine differences among the coordinate models and does not necessarily indicate inconsistent predictions. We consequently focused on atoms with positional SDs below approximately 0.3 Å, whose coordinates were essentially conserved across the datasets. Within this group, atoms with high aBCC_pred_ values generally showed small SDs, whereas lower-scoring atoms showed greater variation. This trend was similar to that observed in the resolution-controlled test datasets (Fig. 4b), indicating that well-supported atomic regions were evaluated consistently across experimental structures of different resolutions.

Together, these analyses distinguish two sources of variation. The Fourier-truncated 5PZL series shows that aBCC_pred_ is largely insensitive to the loss of high-resolution reflections alone, whereas the independently determined 23E structures test whether the learned relationship remains applicable in the presence of additional variation found in experimental crystallographic data. The broadly conserved aBCC_pred_ profiles indicate that the predictions remain comparatively stable under these conditions, although the modest residual differences show that dataset-specific effects are not completely removed. These findings are consistent with our training-data design, in which the original B factors were retained to isolate the information loss caused by the resolution cutoff rather than to reproduce all characteristics of lower-resolution experimental data.

### 3.2. Comparison with experimental omit-map aBCC_act_ and EDIA using 23E dataset

To distinguish the effect of resolution standardization by the prediction model from the properties of the aBCC formulation itself, we calculated aBCC_act_ directly from the experimental omit maps used as inputs to the prediction model for the 23E multiple-resolution dataset (Supplementary Fig. 9a). The experimental-map aBCC_act_ values exhibited atom-wise patterns similar to those of aBCC_pred_. However, the values decreased systematically toward lower resolution, and the spread of the atomic scores progressively narrowed, indicating a reduced ability to distinguish differences among individual atoms.

A similar resolution dependence was observed for EDIA, which decreased systematically toward lower resolution across the ligand, accompanied by a progressive reduction in atom-to-atom score variation (Supplementary Fig. 9b). This behavior is consistent with the formulation of EDIA, in which the evaluation radius is adjusted according to resolution. In addition, EDIA is calculated from an electron-density map used for model building rather than from an omit map and may therefore be influenced by multiple factors, including resolution, atomic displacement, model bias, and coordinate accuracy. Whereas EDIA directly evaluates local support from the experimental electron density, aBCC_pred_ estimates coordinate–density consistency on the scale defined by the standardized 1.5 Å reference framework. The present comparison is therefore intended to examine the resolution dependence of the metrics rather than to compare their ability to classify coordinate errors.

In contrast to aBCC_act_ and EDIA calculated directly from the experimental maps, aBCC_pred_ showed only a limited change across resolutions (Supplementary Fig. 8a). This behavior is consistent with the training design, in which maps spanning different effective resolutions were associated with aBCC values evaluated using a common 1.5 Å reference framework. Thus, aBCC_pred_ reduces the direct dependence of atomic scores on map resolution while retaining variations associated with local coordinate–density consistency and other dataset-specific properties.

### 3.3. Resolution-dependent behavior of aBCC_pred_ in representative cofactors

To examine the robustness of aBCC_pred_ to resolution in comparison with conventional real-space metrics, RSCC values were calculated for structures containing the ubiquitous cofactors SAM, FMN, and ATP across different resolution bins (Fig. 5). Because the chemical structures and binding modes of these cofactors are well established, gross modeling errors are less likely than for newly identified ligands, making them useful test cases for examining the effects of resolution on validation metrics. If aBCC successfully provides a resolution-standardized measure of local coordinate–density consistency, aBCC_pred_ would be expected to show less systematic variation across resolution bins than conventional metrics. It is important to note that aBCC evaluates local density consistency at the atomic level, whereas RSCC is a ligand-level metric reflecting overall shape similarity.

**Figure 5.**
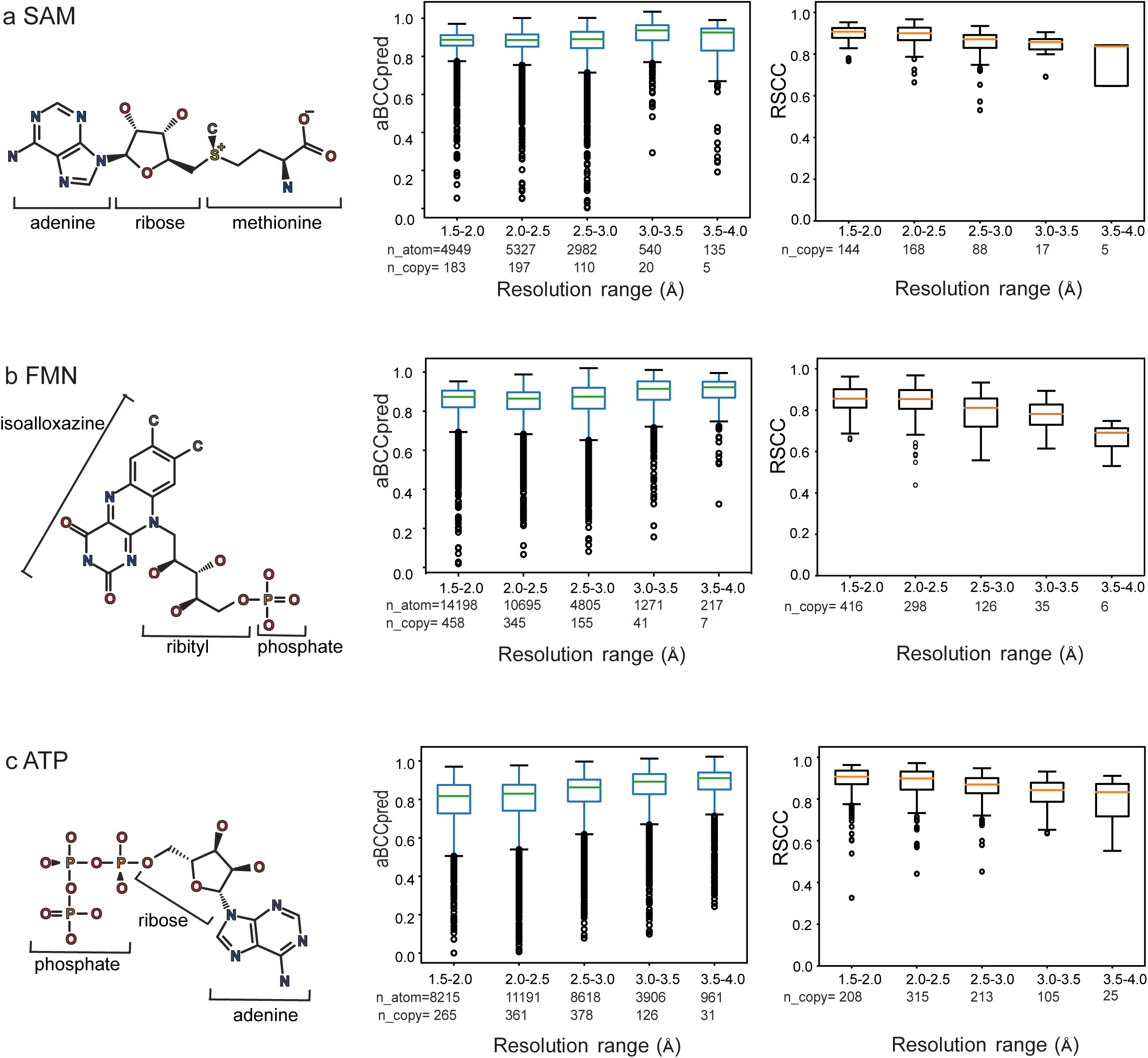
Resolution dependence of aBCC_pred_ and RSCC for representative cofactors. Box plots showing the aBCC_pred_ (left) and RSCC (right) values across resolution bins for SAM (a), FMN (b), and ATP (c). For each cofactor, aBCC_pred_ showed a weaker dependence on resolution than RSCC, indicating a more consistent evaluation of the ligand coordinate–density agreement across a wide resolution range. Box plots are presented to illustrate the overall trends, rather than to support statistical comparisons between the resolution bins.

Consistent with previous observations, ligand-level RSCC values exhibited a systematic decrease as resolution decreased, reflecting the progressive loss of global shape information in real-space correlation metrics. The aBCC_pred_ values showed more nuanced behavior. In contrast to the behavior of RSCC, the overall aBCC_pred_ values remained largely stable across resolution bins. However, a modest increase was observed at a lower resolution, with the magnitude of this effect differing among the cofactors. Specifically, the resolution-dependent increase in aBCC_pred_ values became more pronounced in the order SAM < FMN < ATP (Supplementary Fig. 10), suggesting that this effect may be associated with specific chemical features shared among these cofactors.

To elucidate the origin of this behavior, the aBCC_pred_ values were analyzed by decomposing each cofactor into chemically defined substructures (Supplementary Fig. 10). The phosphate groups exhibited the largest resolution-dependent increase in aBCC_pred_ values. Smaller increases were also observed for certain non-phosphorylated flexible moieties, including the ribose group of ATP, whereas the adenine and isoalloxazine moieties showed only weak or negligible changes. In contrast, no comparable resolution dependence was observed for the methionine moiety of SAM, indicating that the dominant effect was associated with phosphate-containing regions rather than with the cofactor molecules as a whole. Representative examples illustrating the aBCC_pred_ values mapped onto the electron density for each cofactor are shown in Supplementary Fig. 11a–c.

To further investigate whether this trend reflects resolution degradation, the same analysis was performed using Fourier-truncated maps derived from a single high-resolution ATP-bound structure (7PLJ) (Supplementary Fig. 11d). In contrast to the statistical analysis across experimentally determined structures, aBCC_pred_ values were reproduced consistently up to 4.0 Å in the Fourier-truncated dataset, indicating that the pronounced increase observed for phosphate-containing cofactors cannot be explained by resolution degradation alone. Although differences in the structural composition of the datasets across resolution ranges may also contribute to this trend, the current model architecture and training data are plausible factors.

The origin of this phosphate-associated behavior remains unclear. However, the current model has two limitations: First, phosphorus atoms are represented within a generic “other atom” input channel rather than by a phosphorus-specific channel, potentially limiting the model’s ability to learn phosphate-specific density features. Second, phosphorus-containing compounds were relatively underrepresented in the training dataset compared with the major atom types (C, N, and O). Therefore, phosphate-containing ligands are an important limitation of the current model. Improving the representation of less abundant atom types in the training data and incorporating element-specific input features are important for further improving QAEmap and its application to a broader range of ligands.

### 3.4. Application of QAEmap for aBCC evaluation

#### 3.4.1. Comparison of obsolete and replaced data in the PDB

To examine whether QAEmap can capture ligand assignment errors arising from inconsistencies between atomic coordinates and electron density, pairs of obsolete and replaced PDB entries were analyzed. Obsolete entries are updated for various reasons, including ligand misassignment, chemical or geometric corrections, and refinement improvements; however, the specific cause of replacement is not systematically annotated in the PDB. Therefore, obsolete–replacement pairs do not necessarily represent uniform improvements in density fitting and should be interpreted with caution. Across the dataset, changes in the aBCC_pred_ values between the obsolete and replacement entries were not uniform; increases, decreases, and negligible changes were observed. This indicates that obsolete–replacement comparisons cannot be used as straightforward benchmarks to assess improvements in ligand–density consistency.

Nevertheless, closer inspection revealed that cases with increased aBCC_pred_ were enriched in revisions involving the reorientation of rigid aromatic or heterocyclic ring systems (Fig. 6a, b; Supplementary Fig. 12a–d). In these examples, incorrect orientations of the planar ring systems or their substituents in the obsolete models resulted in local mismatches with electron density, whereas the replacement structures exhibited improved consistency, as reflected by the higher aBCC values. Representative ligands included QIH, OF6, ACO, and EA7, although not all cases showed uniform behavior. Notably, aBCC_pred_ is sensitive to subtle coordinate differences that are not readily apparent upon visual inspection. For example, in the BY2-containing structures (Supplementary Fig. 12c), only minor positional differences were observed between the obsolete and replacement models, yet a measurable increase in aBCC_pred_ was detected. This suggests that aBCC_pred_ can capture subtle local improvements in density–coordinate consistency that may not be visually obvious.

**Figure 6.**
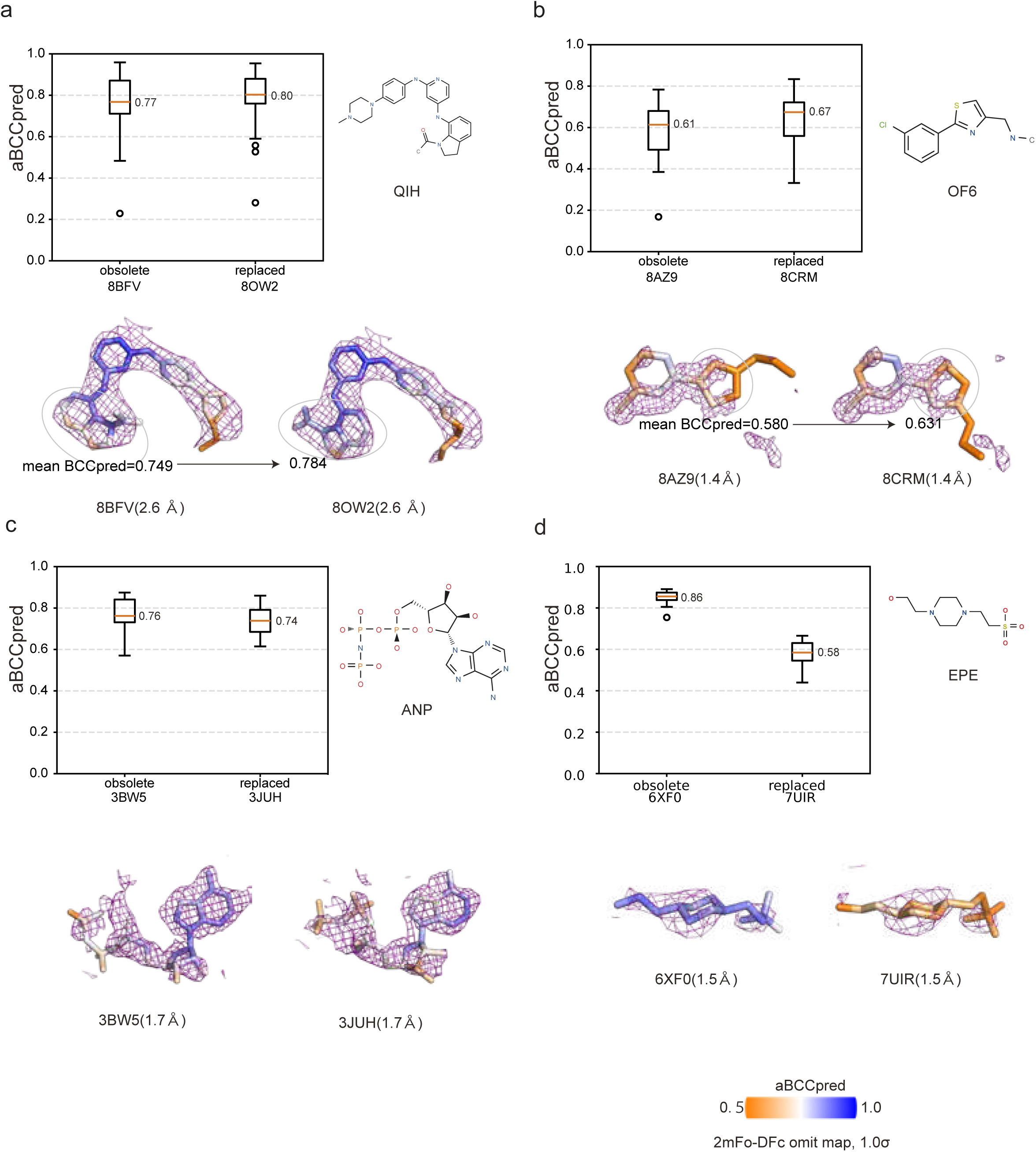
Evaluation of aBCC_pred_ performance in obsolete–replacement structures. (a–d) Examples of obsolete and replaced PDB entries showing cases in which the aBCC_pred_ values increased (a, b), showed limited change (c), or decreased (d) following structural replacement. For each case, box plots summarize aBCC_pred_ values for the obsolete and replaced structures, together with representative electron density maps (2 mFo–DFc, contoured at 1σ). In panels (a) and (b), the mean aBCC_pred_ values calculated for selected aromatic substructures (circled) highlight local improvements in coordinate–density consistency within rigid ring systems. This comparison illustrates the response of aBCC_pred_ to a revision of the deposited atomic model and does not imply that the molecular identity or coordinates in the replacement entry are necessarily correct.

Conversely, some cases did not show substantial changes in aBCC_pred_ despite the apparent differences in ligand orientation. For instance, in RTX-containing structures (Supplementary Fig. 12d), only marginal differences in aBCC_pred_ were observed between the obsolete and replacement entries. This highlights the limitations associated with the sensitivity of the metric or the ambiguity in the underlying electron density. In contrast, revisions involving more flexible ligands—such as those dominated by aliphatic chains or phosphate-containing groups—showed little or inconsistent changes in aBCC_pred_ (Fig. 6c,d). In these cases, the electron density was comparatively diffused and did not strongly constrain the ligand geometry, thereby limiting the sensitivity of aBCC_pred_ to coordinate differences.

This behavior is consistent with the observations from the cofactor analysis. In some cases, differences in aBCC_pred_ were observed even when the underlying electron density remained poorly defined, complicating the interpretation. For example, for the buffer molecule EPE (Fig. 6d), substantial differences in aBCC_pred_ were observed between the obsolete and replacement entries, despite both models being associated with weak or ambiguous electron densities. Because the electron density maps are derived independently from the deposited structure factors for each entry, such differences may reflect not only coordinate changes but also variations in map quality, and therefore cannot be unambiguously interpreted as improvements in ligand assignment.

Together, these observations indicate that aBCC_pred_ is the particularly informative for detecting local coordinate discrepancies in structurally constrained regions, such as rigid aromatic or heterocyclic systems, whereas its sensitivity is reduced for flexible ligands or poorly defined electron densities. These findings highlight both the applicability and limitations of aBCC_pred_, supporting its role as a complementary locally focused measure of coordinate–density consistency.

## 4. Conclusions

In this study, we present a framework for evaluating the consistency between ligand atomic coordinates and electron density in macromolecular structures using a machine learning model trained on standardized reference maps. Rather than treating coordinate reliability as being uniformly constrained by global resolution, QAEmap evaluates local coordinate–density consistency against a common high-resolution reference framework. This approach enables resolution-standardized assessment at the individual atom level, offering a new perspective that complements conventional metrics, such as RSCC and EDIA.

This framework is intended to support the interpretation of ligand structures rather than provide a binary assessment of model correctness. The purpose of aBCC is not to classify ligand models as correct/incorrect but to quantify the local coordinate–density consistency of atomic coordinates based on standardized electron density. We anticipate that this information will complement, rather than replace, the judgment of crystallographers by communicating atom-wise coordinate confidence to downstream users in structure-based drug design.

Although the aBCC is defined as a resolution-standardized metric, the prediction accuracy decreases at lower resolutions because the information content of the input density maps is inherently reduced. Further improvements in the model architecture and training data may extend the applicability of this framework to lower-resolution structures.

Our analyses demonstrated that aBCC is particularly sensitive to coordinate discrepancies in structurally constrained ligand moieties, such as aromatic and heterocyclic ring systems. In contrast, the performance was more variable for flexible ligand regions and phosphate-containing moieties, highlighting areas for future model improvement. By standardizing the density evaluation across different resolutions, this approach establishes a foundation for the systematic and scalable assessment of local coordinate reliability in macromolecular structures. Such atom-wise confidence information may help identify ligand regions that warrant further computational exploration or cautious interpretation.

Future developments include the incorporation of richer atomic descriptors, expansion of the training dataset, and integration of the framework into practical workflows. Overall, our findings demonstrate the feasibility and utility of learning-based atom-resolved metrics for evaluating coordinate–density consistency and provide a basis for further methodological advances toward more reliable structural interpretation in structural biology and structure-based drug discovery.

## Acknowledgements

We thank Prof. Yasushi Okuno for establishing the LINC consortium and enabling access to the computational infrastructure. We are grateful to Dr. Keitaro Yamashita for guidance on the use of Servalcat in Fc map construction. We thank Prof. Genji Kurisu and Dr. Gert-Jan Bekker for valuable discussions on the development and future applications of this study. We thank MOLSIS Inc. for their support with the docking calculations. We also thank ChatGPT (OpenAI) for assistance in technical scripting. This research was supported by the Platform Project for Supporting Drug Discovery and Life Science Research (Basis for Supporting Innovative Drug Discovery and Life Science Research (BINDS)) from AMED under grant number JP26ama121023 (M.I.). This research used computational resources provided by Yokohama City University.

## Data availability

All crystallographic structures used to construct the training and evaluation datasets in this study are publicly available in the Protein Data Bank (https://www.rcsb.org) under the PDB accession codes listed in the Methods (Section 2) and Supplementary Information. No new macromolecular structures were determined in this work. Derived data supporting the findings of this study, including the training and test datasets, simulated resolution-series maps, docking poses, and example density boxes, are available from the corresponding author upon reasonable request.

## Code availability

The QAEmap program is freely available for academic use through GitLab (https://gitlab.com/qaemap_products).

## Software

Docking poses were generated using the Molecular Operating Environment (MOE v. 2022.02; Chemical Computing Group ULC, Montreal, QC, Canada). The software package PyMOL (The PyMOL Molecular Graphics System, v. 2.3, Schrödinger, LLC, https://www.pymol.org/2/) was used for the visualization of protein structures and maps. CCP4 (v. 7.0, Collaborative Computational Project No. 4, https://www.ccp4.ac.uk/) and Servalcat were used for creating protein structures and map files. Tensorflow (v. 1.x, Google Brain, https://www.tensorflow.org/install/pip) and Python (v. 3.6, Python Software Foundation, https://www.python.org/downloads/) were used for developing QAEmap.

---

**Fig. 7.**
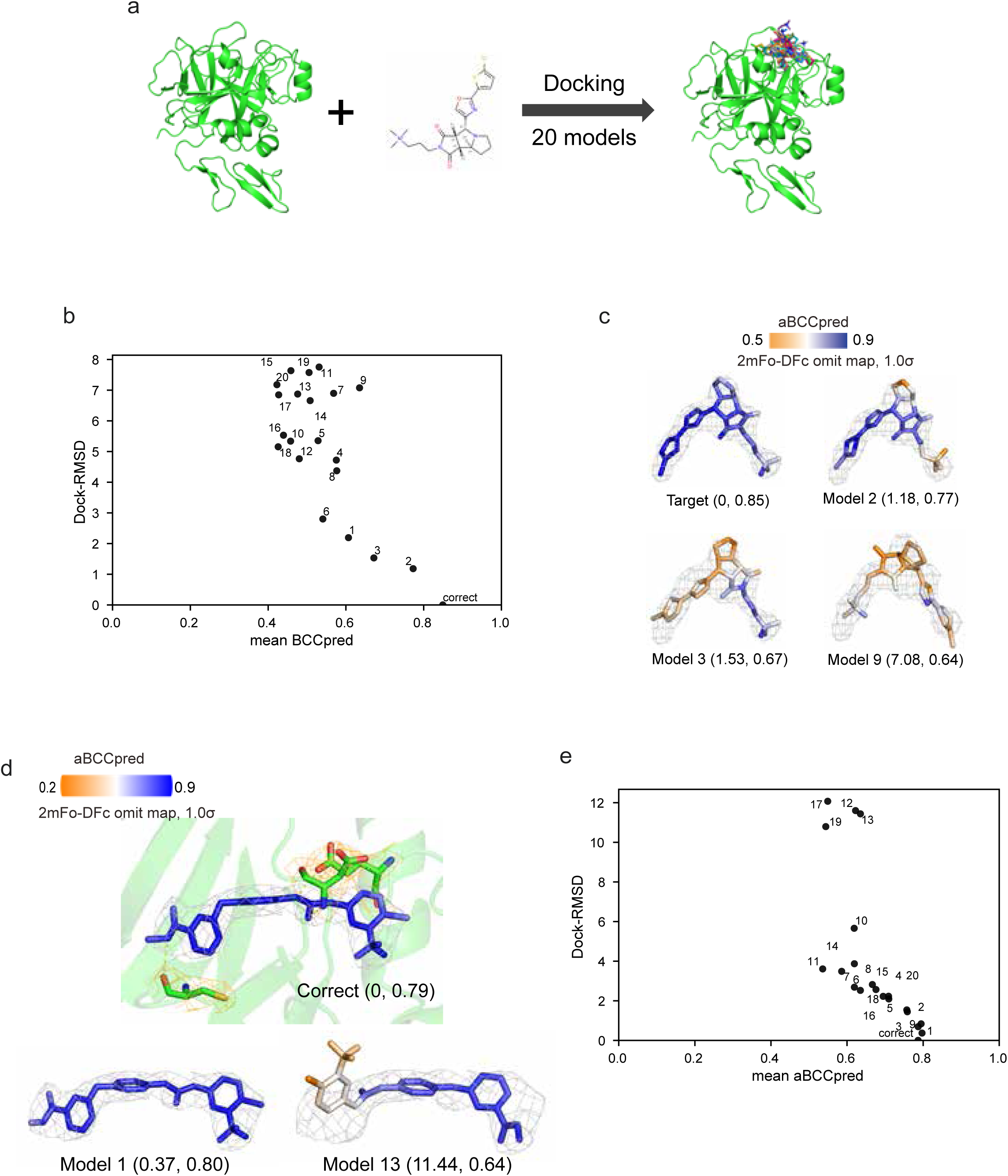

**Figure S1.**
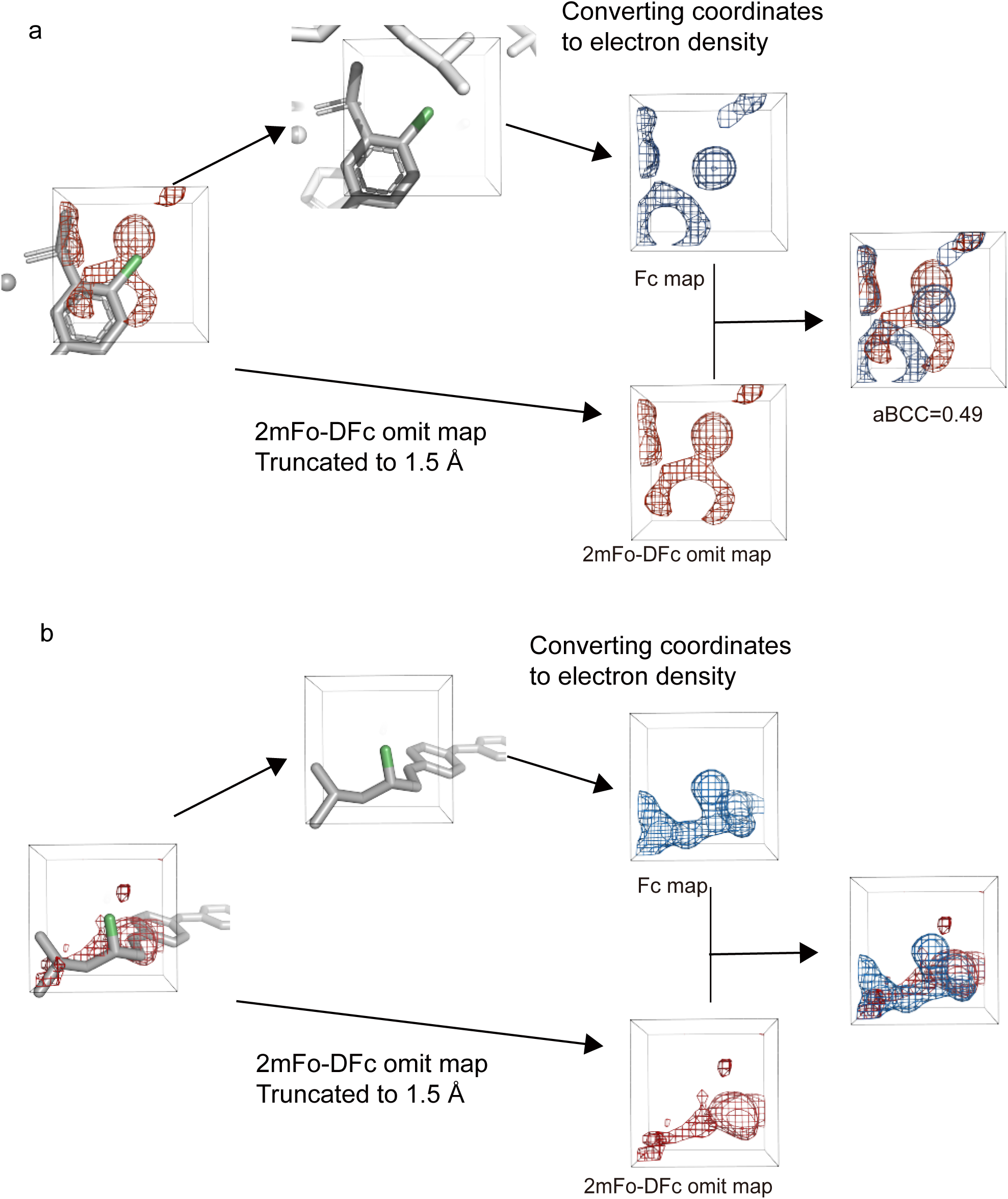
Examples of low aBCC values caused by incorrect or uncertain atomic coordinates. (a) Example of incorrect ligand conformation. The atomic coordinates deviated from the correct conformation, resulting in a mismatch between the Fc map (blue) and 2 mFo**–**DFc omit map (red), yielding a low aBCC score (0.49). (b) Example of a flexible ligand region with high B-factor. Although the atomic coordinates were geometrically reasonable, the 2 mFo–DFc omit map showed a weak or ambiguous density. This resulted in poor consistency with the calculated Fc map and a low aBCC score (0.55). Together, these cases illustrate that the aBCC values decrease when the coordinates are incorrect or poorly supported by the experimental data, even after refinement.

**Figure S2.**
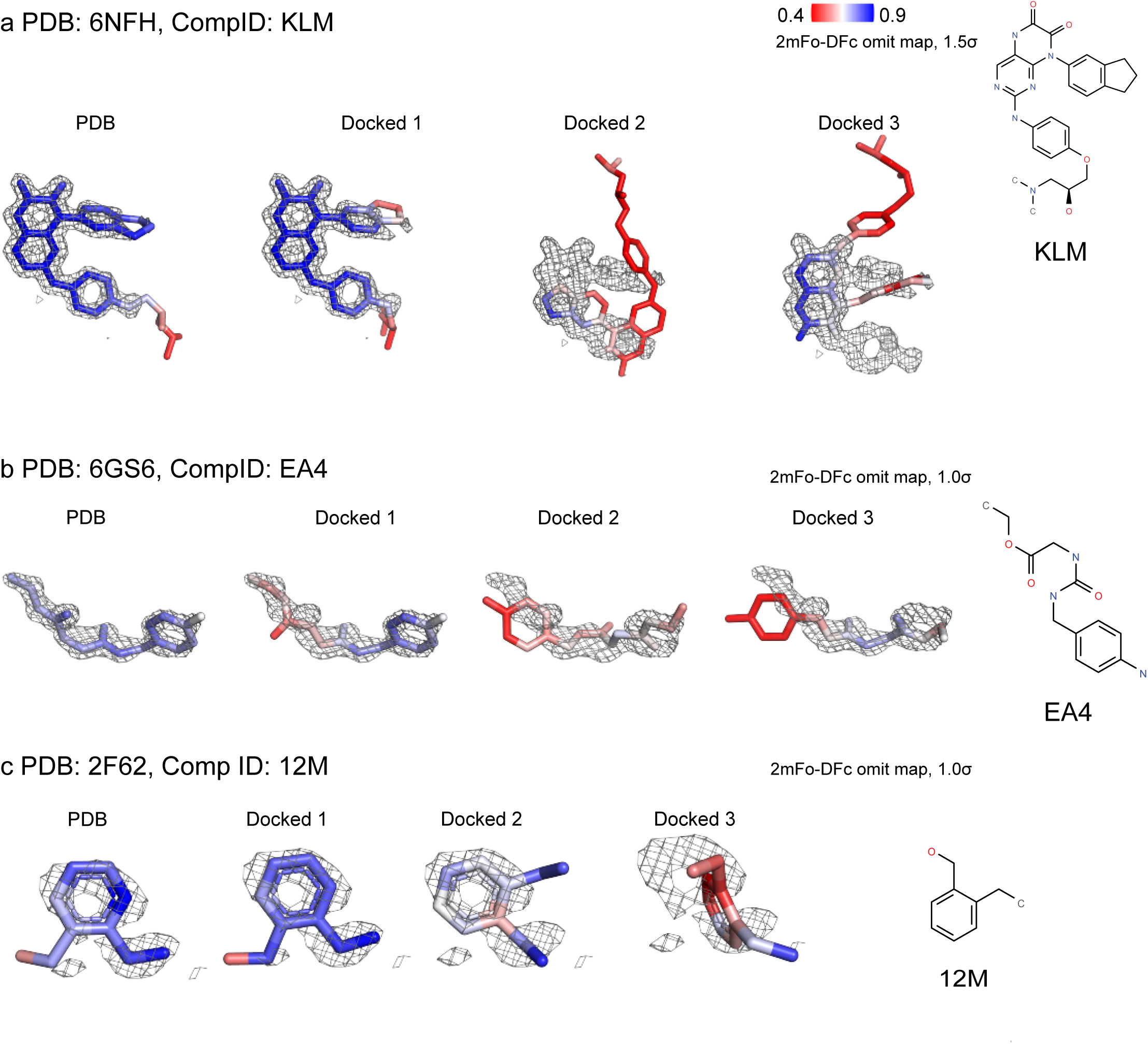
Representative aBCC_act_ values for ligands in deposited models and docking poses. For each compound, ligands from the deposited models together with three docking poses (Docked 1–3) generated by MOE are shown. The aBCC_act_ values were mapped onto each ligand to visualize the consistency between the atomic coordinates and electron density. Incorrect or poorly fitted poses exhibited locally reduced aBCC_act_ values relative to the PDB reference pose. Chemical structures are shown on the right for reference.

**Figure S3.**
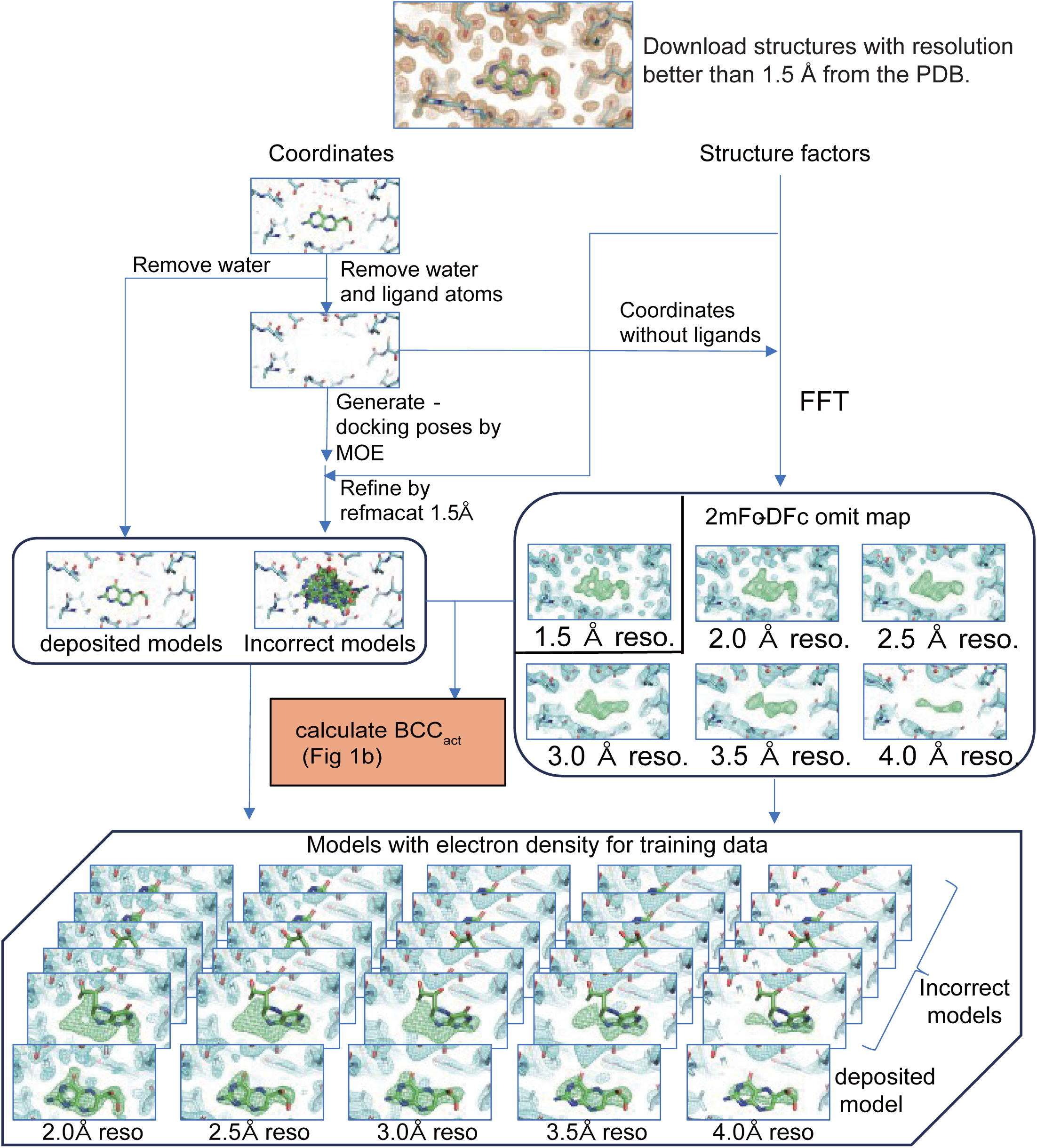
Workflow for generating training dataset. Structures with resolutions of 1.5 Å or better were downloaded from the PDB. For each complex, deposited models and up to 20 docking poses generated using MOE were prepared. Water molecules and ligand atoms were removed as appropriate, and all models were refined at 1.5 Å resolution using REFMAC. For each model, 2 mFo–DFc omit maps were generated at multiple resolutions (2.0–4.0 Å). The aBCC_act_ values were calculated by comparing the refined Fc map with the corresponding omit map at 1.5 Å resolution. The coordinates and omit maps at lower resolutions (2.0–4.0 Å) were used as input features, while the corresponding aBCC_act_ values served as training targets.

**Figure S4.**
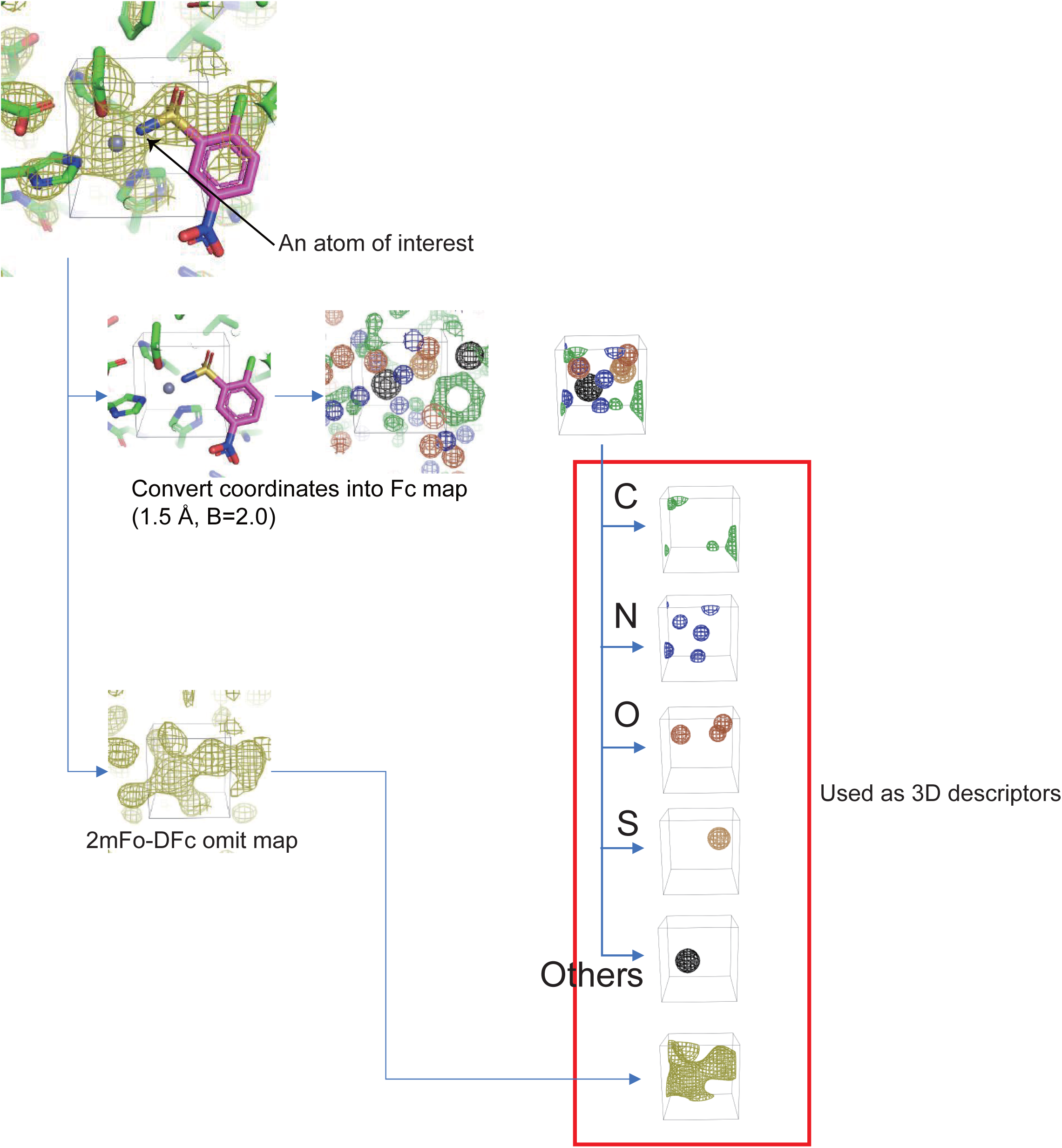
Construction of atom-level 3D descriptors for training. For each atom of interest, a local cubic box centered on the atom was extracted from two sources: an Fc map calculated from the refined coordinates at 1.5DÅ resolution with a uniform B-factor of 2.0 **Å²**, separated into atom-type-specific channels (C, N, O, S, and others); and the 2 mFo–DFc omit map at the corresponding resolution (2.0–4.0DÅ). These components were combined to form multichannel 3D descriptors that serve as input features for model training.

**Figure S5.**
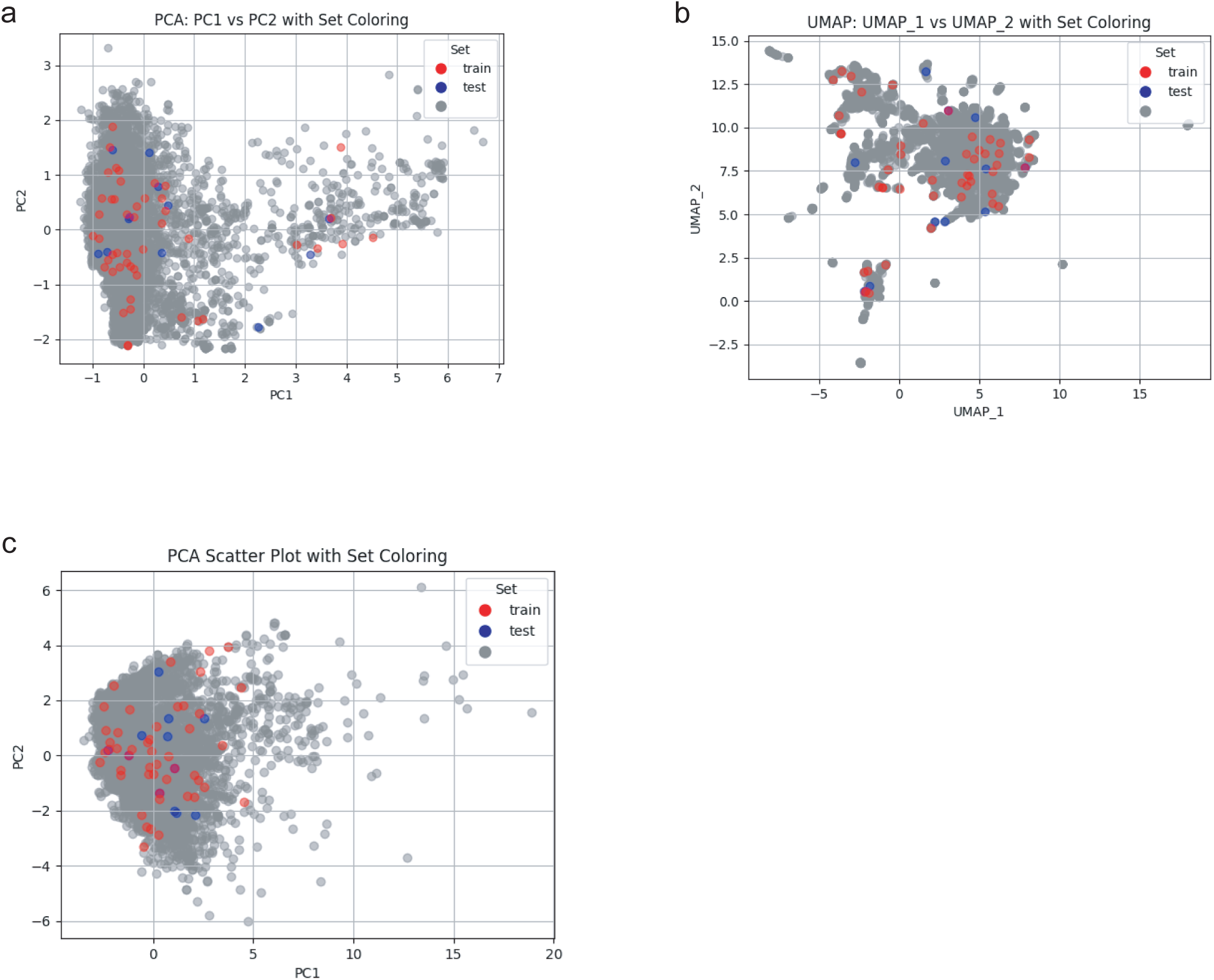
Chemical space distribution of compounds used in this study. The chemical space of the compounds was visualized using principal component analysis (PCA) and uniform manifold approximation, and projection (UMAP). Molecular representations were generated using either ECFP4 fingerprints (2048 bits) or a set of physicochemical descriptors (SlogP, AMW, NumRotatableBonds, NumRings, NumAromaticRings, NumHBA, NumHBD, TPSA, and CSP3). The compounds are colored according to their assignment to the training or test sets. (a) PCA projection based on ECFP4 fingerprints illustrating the structural diversity of the compounds. (b) UMAP based on ECFP4 fingerprints highlighting local similarity relationships among compounds. (c) PCA projection based on physicochemical descriptors illustrating the coverage of the physicochemical property space. Across all the projections, the training set broadly covers the chemical space occupied by the test set, indicating that the test compounds lie largely within the domain represented by the training data.

**Figure S6.**
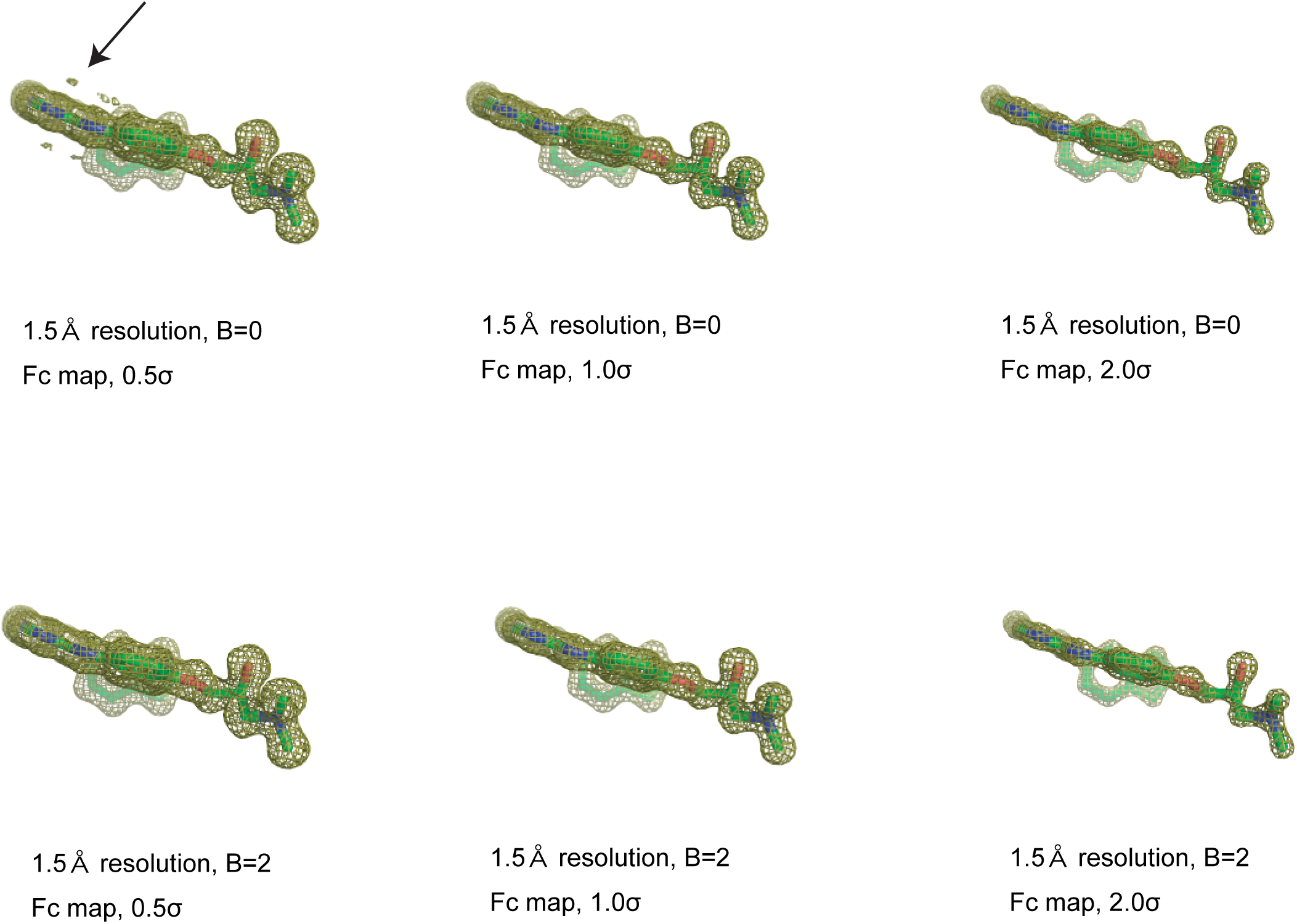
Effect of B-factor on Gibbs artifacts in Fc maps at 1.5 Å resolution. Fc maps calculated at 1.5 Å resolution are shown for two different B-factor settings, B = 0 **Å²** (top row) and B = 2.0 **Å²** (bottom row), contoured at 0.5σ, 1.0σ, and 2.0σ. Strong oscillatory features characteristic of Gibbs artifacts are observed at B = 0 **Å²** (arrow), whereas these artifacts are markedly reduced when a finite B-factor (B = 2.0 **Å²**) is applied.

**Figure S7.**
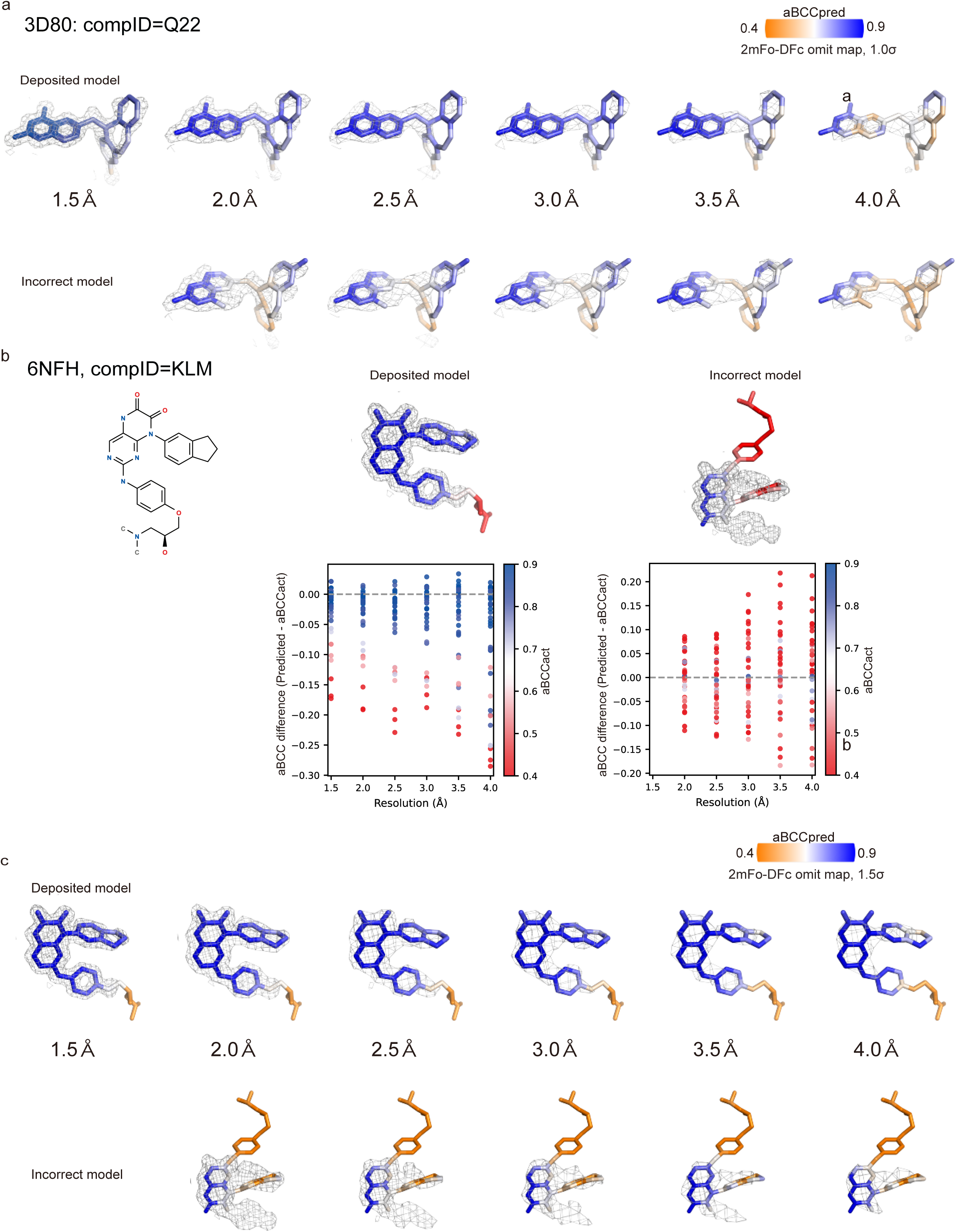
Representative visualization of aBCC prediction across resolutions. **(a)** Representative correct and incorrect ligand models of 3D80 colored according to aBCC_pred_ values (orange–white–blue). The corresponding prediction errors relative to the reference aBCC_act_ values are summarized in Fig. 4b. **(b)** Prediction errors relative to reference aBCC_act_ values for representative correct and incorrect ligand models of 6NFH. **(c)** Corresponding ligand structures of 6NFH colored according to the predicted aBCC_pred_ values (orange–white–blue). For all examples, atomic coordinates were kept identical, and only the map resolution was varied (1.5–4.0 Å). Electron density is shown as 2 mFo–DFc omit maps contoured at 0.5σ. Although the prediction errors gradually increased at lower resolutions, the spatial distribution of the predicted aBCC values remained largely unchanged, indicating consistent identification of high- and low-coordinate–density consistency atomic regions across the tested resolution range.

**Figure S8.**
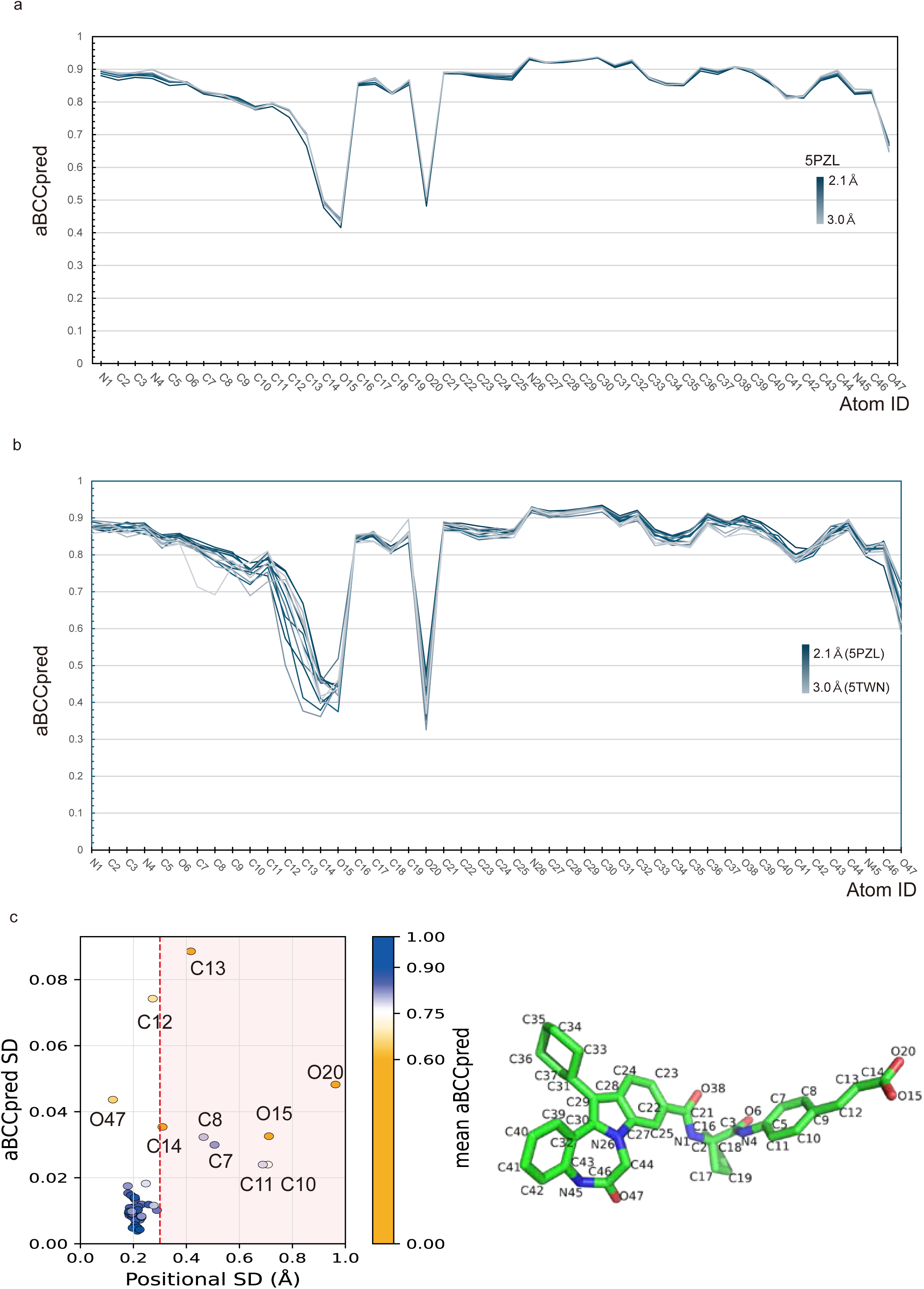
Resolution robustness and atom-wise consistency of aBCC_pred_ for ligand 23E. (a) Atom-wise aBCC_pred_ profiles calculated from the original 2.1 Å diffraction data for 5PZL and from maps generated by Fourier truncation of the same data to resolutions of 2.3–3.0 Å. Line colors indicate resolution, progressing from dark blue for the original 2.1 Å data to light gray for the 3.0 Å truncated data. The profiles remained nearly unchanged after truncation, with a difference of approximately 0.01 in mean aBCC_pred_ over the resolution range examined. (b) Atom-wise aBCC_pred_ profiles for ligand 23E in 13 independently determined structures of the same protein and crystal form, with resolutions ranging from 2.1 to 3.0 Å. Line colors indicate resolution, progressing from dark blue for the highest-resolution dataset (5PZL, 2.1 Å) to light gray for the lowest-resolution dataset (5TWN, 3.0 Å). The overall profiles were broadly conserved across the datasets, although notable variation was observed for a limited set of atoms, including C10–C14, O15, and O20. (c) Relationship between the positional SD and the SD of aBCC_pred_ for each atom across the 13 independently determined structures. Each point represents one ligand atom and is colored according to its mean aBCC_pred_ across the datasets. The vertical dashed line indicates a positional SD of 0.3 Å. Among atoms with positional SDs below this threshold, high-scoring atoms generally showed small variation in aBCC_pred_, whereas lower-scoring atoms showed greater variation. Atoms with larger positional SDs may reflect differences in ligand conformation among the independently refined structures. Selected atoms are labeled, and the chemical structure on the right shows the atom numbering used in the plots.

**Figure S9.**
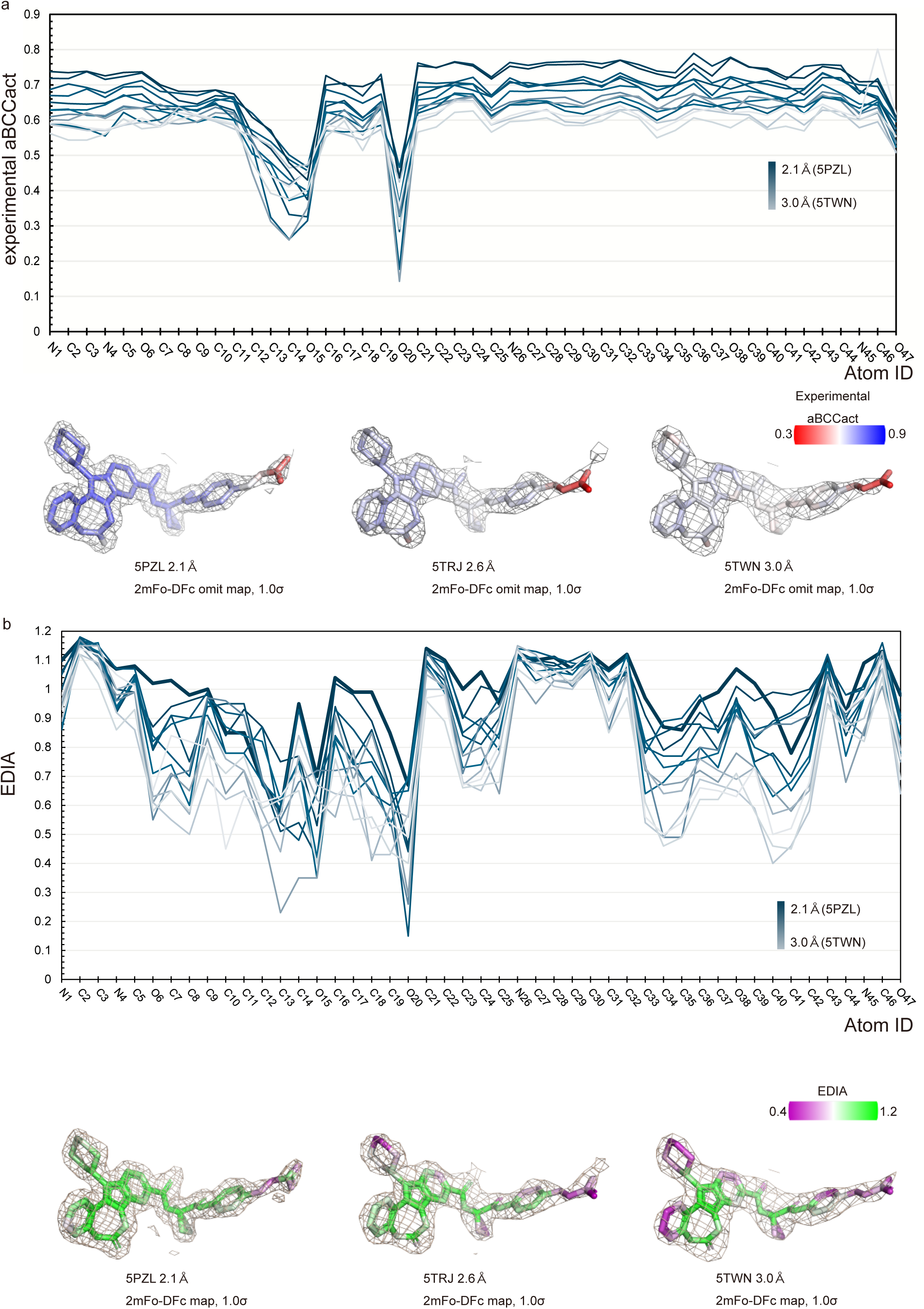
Resolution dependence of experimental omit-map aBCC_act_ and atomic EDIA values for multiresolution 23E dataset. (a) Experimental omit-map aBCC_act_ values for the representative structures shown in Fig. 4c. Each line corresponds to a single structure, with colors ranging from dark to light blue as the resolution decreases. Representative ligand electron density maps (2 mFo–DFc omit maps, contoured at 1.0σ) are shown below. The experimental omit-map aBCC_act_ values decrease systematically with decreasing resolution, indicating their dependence on map resolution. (b) Atomic EDIA values for same structure. Colors are ordered identically to panel (a) from high to low resolution. Representative ligand models are colored according to the atomic EDIA values. Similar to the experimental omit-map aBCCact, the experimental omit-map aBCC_act_, the atomic EDIA values exhibited a clear resolution dependence, demonstrating that both metrics were influenced by the map resolution.

**Figure S10.**
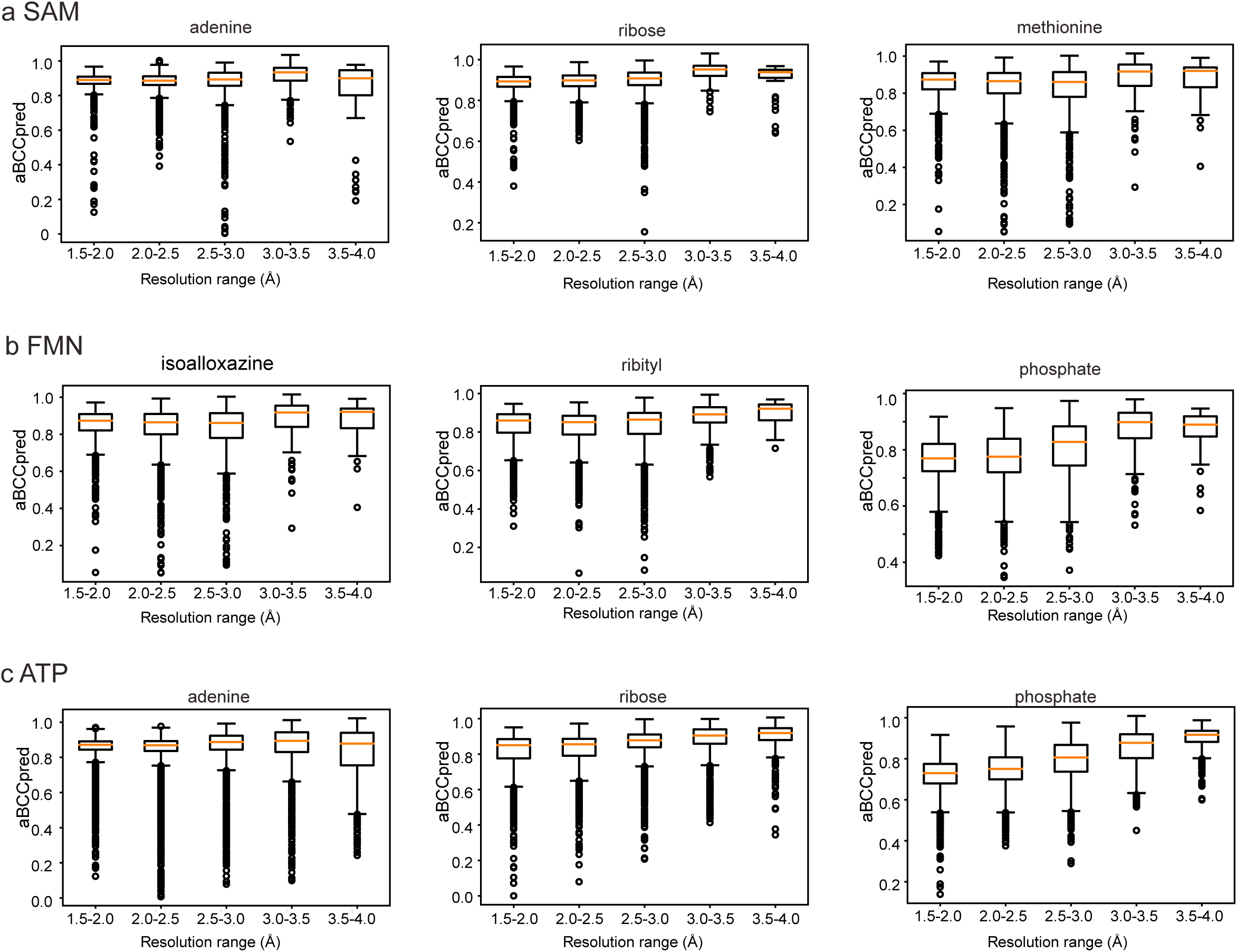
Resolution dependence of aBCC_pred_ on cofactor substructures. Cofactors were decomposed into chemically distinct substructures, and aBCC_pred_ values were analyzed for each substructure across resolution ranges (1.5**–**2.0, 2.0**–**2.5, 2.5**–**3.0, 3.0**–**3.5, and 3.5**–**4.0 Å). The box plots represent the distribution of aBCC_pred_ values within each substructure and resolution bin. (a) SAM, (b) FMN, and (c) ATP. The substructures were defined as adenine, ribose, and methionine for SAM; isoalloxazine, ribityl, and phosphate for FMN; and adenine, ribose, and phosphate for ATP.

**Figure S11.**
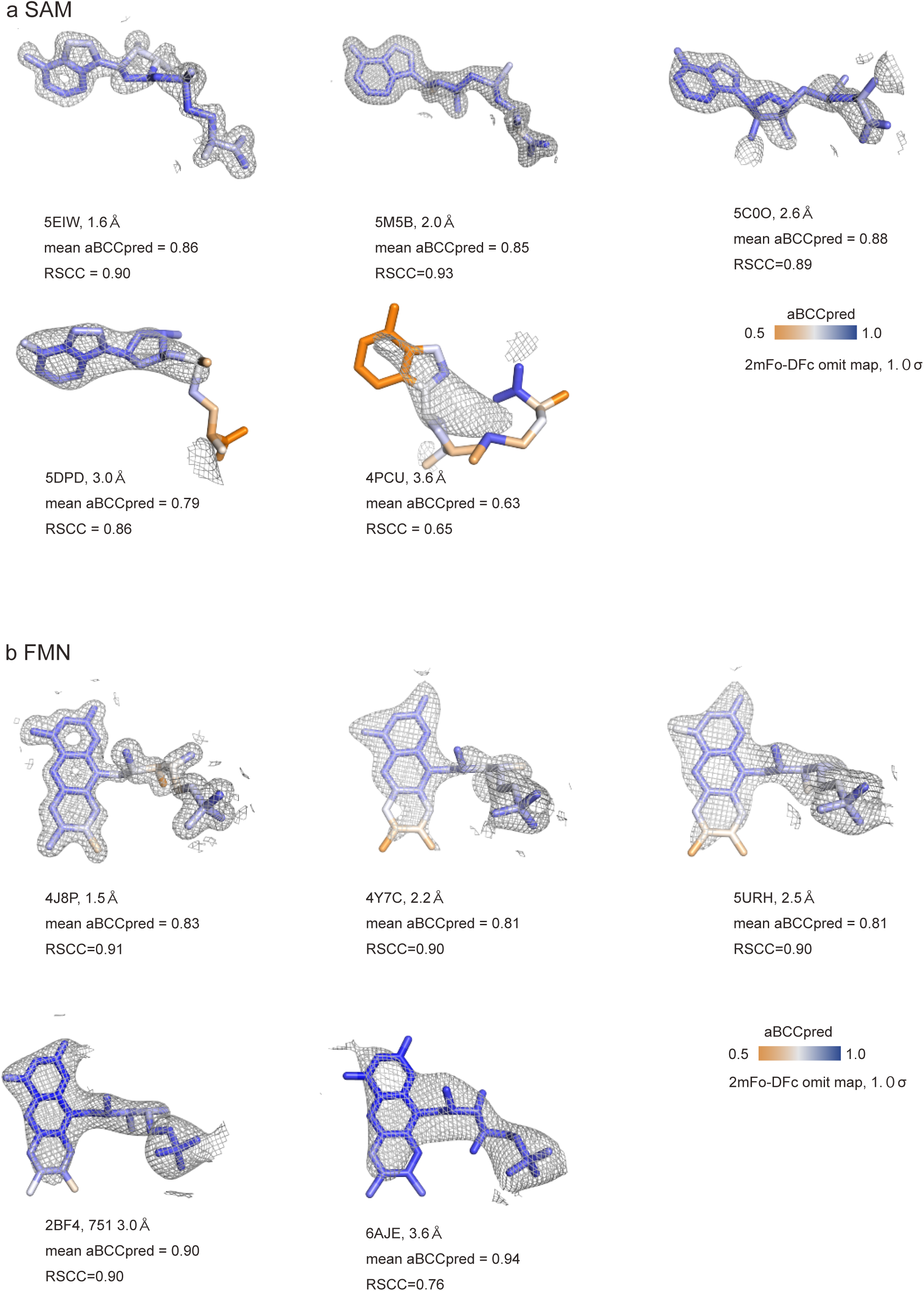
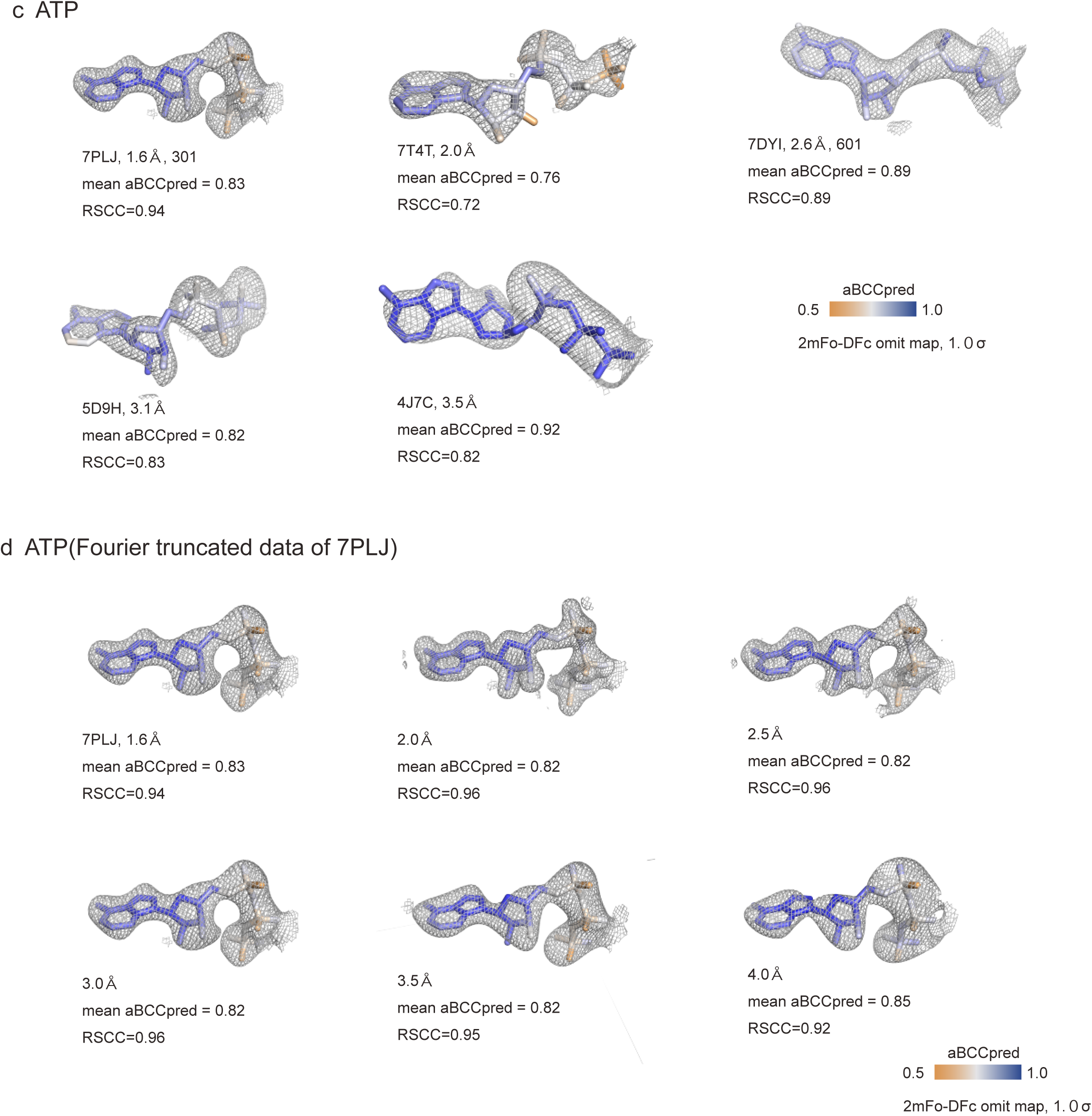
Representative examples of aBCC_pred_ and RSCC values mapped onto electron densities for cofactors. Electron density maps (2 mFo–DFc omit maps, contoured at 1σ) are shown for representative structures containing (a) SAM, (b) FMN, and (c) ATP across different resolution ranges. Atomic coordinates are colored according to the aBCC_pred_ values, and the mean aBCC_pred_ values averaged over the ligand atoms are shown for each structure. Corresponding ligand-level RSCC values for each structure are shown. (d) Control analysis using Fourier-truncated maps derived from a single high-resolution ATP-bound structure (PDB ID: 7PLJ). Structure factors were truncated to lower resolutions (2.0–4.0 Å), and maps were calculated under identical conditions to assess the effects of resolution cutoff while minimizing structural variability. These examples demonstrate that where RSCC decreases with decreasing resolution, aBCC_pred_ values remain relatively stable and reflect local density features, with modest increases observed in the phosphate groups at lower resolutions.

**Figure S12.**
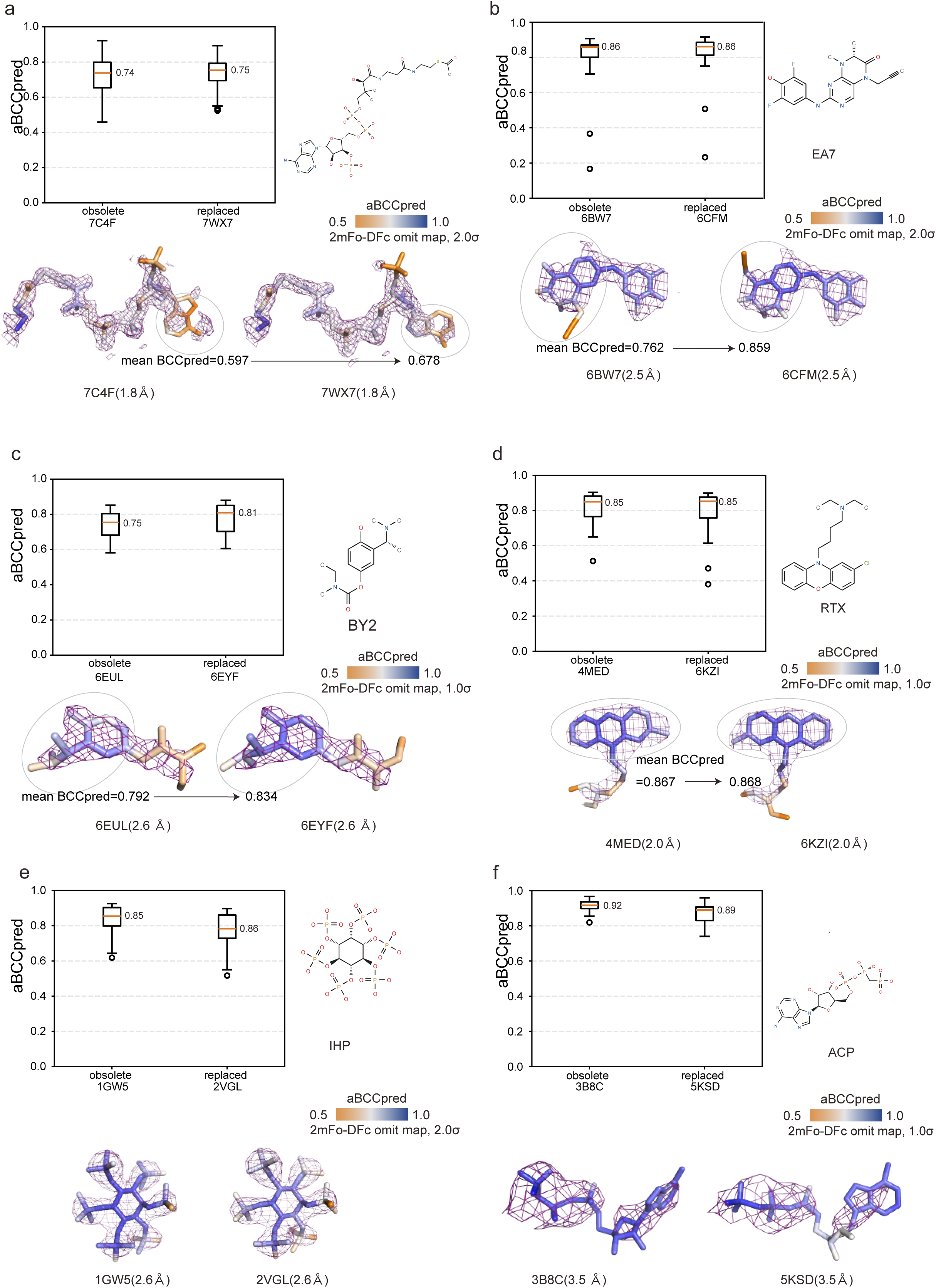
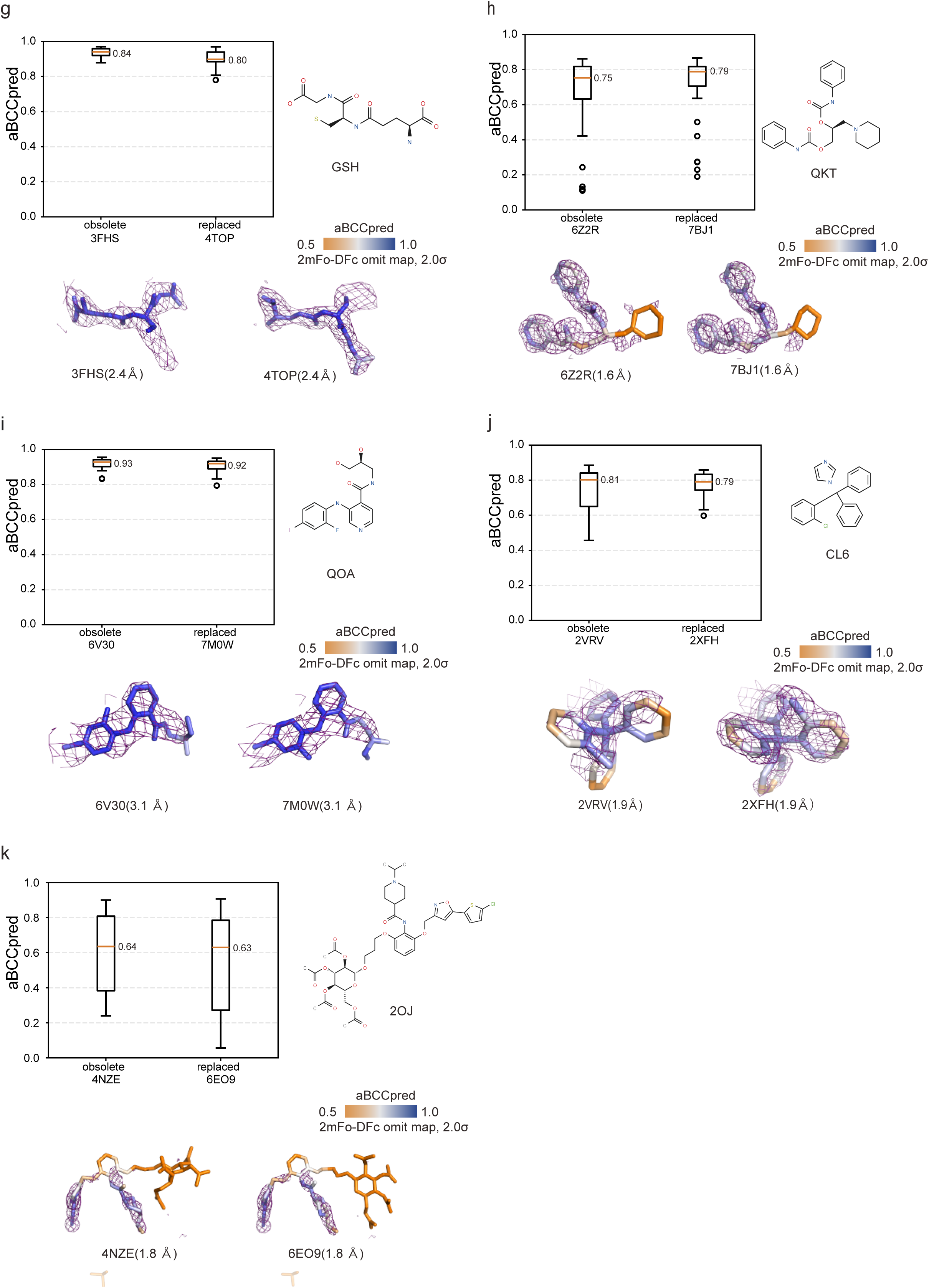
Additional examples of obsolete and replaced PDB entries analyzed using aBCC_pred_. (a–d) Cases involving corrections of aromatic or heterocyclic ring systems. In panels (a–c), the structural revisions are accompanied by clear increases in aBCC_pred_, indicating improved consistency between the atomic coordinates and electron density. By contrast, panel (d) represents a case in which the revision does not lead to a clear change in aBCC_pred_, suggesting a limited impact on local density consistency. The mean aBCC_pred_ values calculated for selected aromatic substructures (circled) are shown to highlight local improvements in coordinate–density consistency within rigid ring systems. (e–k) Examples of modifications other than aromatic ring corrections. In these cases, changes in aBCC_pred_ were generally modest, indicating that not all coordinate revisions were associated with detectable differences in local density consistency. Ligands are displayed as sticks colored according to aBCC_pred_ values, and electron density is shown as 2 mFo–DFc omit maps contoured at 2σ.

